# Episodic memory facilitates flexible decision making via access to detailed events

**DOI:** 10.1101/2025.03.13.643066

**Authors:** Jonathan Nicholas, Marcelo G. Mattar

**Affiliations:** Department of Psychology New York University New York, NY 10003

**Keywords:** decision making, episodic memory, incremental learning

## Abstract

Our experiences contain countless details that may be important in the future, yet we rarely know which will matter and which won’t. This uncertainty poses a difficult challenge for adaptive decision making, as failing to preserve relevant information can prevent us from making good choices later on. One solution to this challenge is to store detailed memories of individual experiences that can be flexibly accessed whenever their details become relevant. By allowing us to store and recall specific events in vivid detail, the human episodic memory system provides exactly this capacity. Yet whether and how this ability supports adaptive behavior is poorly understood. Here we aimed to determine whether people use detailed episodic memories to make decisions when future task demands are uncertain. We hypothesized that the episodic memory system’s ability to store events in great detail allows us to reference any of these details if they later become relevant. We tested this hypothesis using a novel decision making task where participants encoded individual events with multiple features and later made decisions based on these features to maximize their earnings. Across five experiments (total n = 535), we found that participants referenced episodic memories during decisions in feature-rich environments, and that they did so specifically when it was unclear at encoding which features would be needed in the future. Overall, these findings reveal a fundamental adaptive function of episodic memory, showing how its rich representational capacity enables flexible decision making under uncertainty.

## 1 Introduction

Humans possess the remarkable capacity to remember individual events from the past in high fidelity using our episodic memory system ^1,2^. From merely a single exposure, we can recall complex experiences consisting of many different details — the taste of a meal, the layout of a restaurant, the conversation around a dinner table — even when it is unclear whether we will actually ever need any of this information again. Why do we maintain such detailed memories of past events?

One possible answer is that this ability is adaptive, because the aspects of an experience that will be relevant for future behavior are rarely apparent when they are first encountered. For instance, when deciding whether to visit a restaurant, you may need to recall details that weren’t previously important during your prior meals there, such as whether the menu can accommodate your vegan friend or if the ambiance is appropriate for a work meeting. By storing detailed representations of past experiences, our episodic memory system can allow us to access any of these details if they ever become relevant in the future. In this way, episodic memory may enable flexible decision making by allowing us to repurpose our memories for novel goals.

Despite a long history of work on the adaptive role of episodic memory ^3,4,5,6^, computational research has only recently started to characterize the advantages it can provide for decision making ^7,8,9^. A key theme in this work is that the episodic memory system is poised to address several shortcomings faced by other forms of memory, particularly incremental learning, which aids decision making by allowing us to gradually estimate the value of different choice options over many repeated encounters ^10,11,12,13,14,15,16^. Importantly, incremental learning suffers from a well-known problem called the *curse of dimensionality*, which arises because computational and memory demands rapidly scale with the richness of what must be learned ^17^. For instance, when learning about choice options with multiple features, an agent capable of only incremental learning would need to track and update values for each independently.

One way humans may circumvent this issue is by selectively learning about only the most relevant features of experience while ignoring information about the rest ^18^. Using selective attention in this way can allow for efficient decisions in high-dimensional environments — when a feature is deemed relevant, its value is tracked and incrementally updated, requiring only a simple retrieval at choice time. Such a strategy is highly effective under circumstances in which feature relevance can be reliably inferred, and people likely employ it when this is the case ^19,20^. But environments like this are, in reality, rare. Further, augmenting incremental learning with selective attention enforces rigidity, ultimately harming future choices that may depend on features that were initially ignored.

There are also fundamental limits to the number of features that can be reasonably attended to and tracked simultaneously ^21^, making this strategy increasingly impractical in the real world. These limitations are precisely the types of problems that episodic memory’s detailed representations are best equipped to address.

A separate but related challenge is that incremental learning is most successful when experiences are repeated, yet actual experiences are unlikely to be encountered more than a single time. Episodic memory, by contrast, allows us to encode and retrieve individual experiences, making it naturally suited to real world environments where experiences rarely repeat ^8^. Indeed, this capacity for one-shot learning has dominated research on episodic memory’s role in decision making, where much work has demonstrated that humans can effectively guide their decisions by retrieving and leveraging individual past experiences ^22,23,24,25,26,27^. Yet, despite this progress, whether the episodic memory system’s ability to encode detailed experiences provides its own advantages for decision making remains largely unknown. Our primary goal was to address this gap.

Here we hypothesized that i) people use episodic memory to access details from past events for decision making in feature-rich environments, and that ii) this strategy enables flexibility when it is unclear which details will be needed in the future. To directly test these ideas, we developed a novel decision making task where participants were asked to encode individual episodes consisting of items that varied across multiple features (e.g., color and category) and an associated reward (**Figure 1A-B**). After encoding these episodes, participants made value-based decisions in which they were shown offers consisting of specific features (e.g., red or animal), and were then asked to either take or leave each offer. In order to determine the value of these offers, participants needed to sum the rewards associated with all items that had the offered feature in common (e.g., all red things).

**Figure 1:**
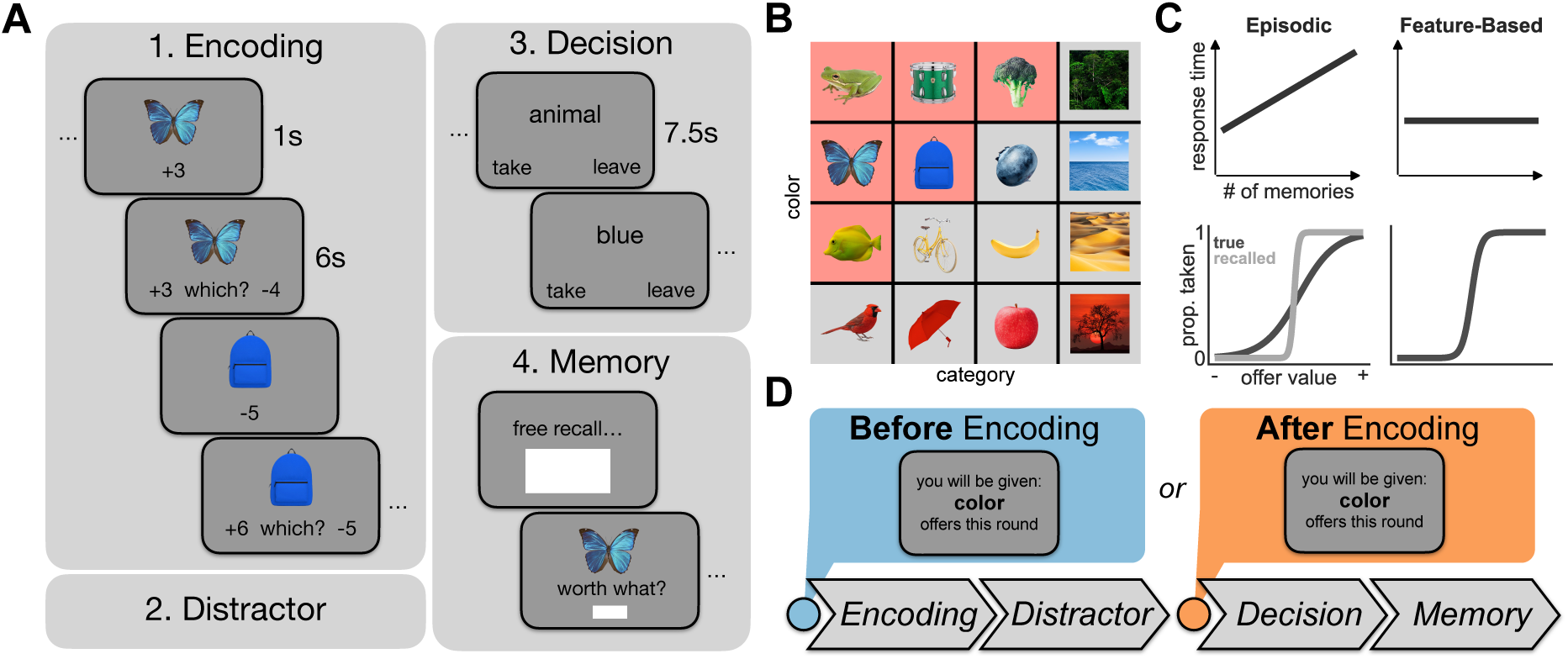
Task Design. **A)** The four phases completed by participants in each round of the task used in Experiments 1-4. In the first part of a round, participants completed an encoding phase which allowed them to encode individual items and an associated reward (an episode). Immediately after viewing the episode, participants completed an attention check consisting of the item alongside two options, either the associated reward that was just shown or another randomly selected reward. Participants completed six trials per round. Following encoding, participants then completed a 90 second distractor task consisting of a 2-back working memory task. This was designed to prevent active rehearsal of the episodes. Following the distractor, participants completed a decision phase in which they were shown offers consisting of a single feature and were asked to either take or leave this offer. The value of each offer was the sum of all episodes described by the offer. Lastly, after the decision phase, participants completed a subsequent memory test consisting of two tasks. They were first asked to freely recall each of the items they had seen in the round. They were then shown each item and asked to provide the reward that was associated with each item. Participants completed between 5 and 9 rounds, depending on the experiment. **B)** The full set of stimuli shown to participants in experiments one, three, and four. Stimuli differed along two dimensions: color and category. A subset of six images were sampled from these to be shown in each round (with an example shown in red). See **Figure S1** for the stimuli used in experiment two. **C)** Simulated choice behavior on the decision phase from two toy process models implementing either an *episodic* or *feature-based* decision strategy (See **Methods** for details). During each offer, the episodic model randomly samples individual items and their associated reward without replacement. To make choices, it then sums up the recalled rewards associated with all recalled offer-relevant items that match the offer, and then chooses to either take or leave each offer depending on whether this decision variable was positive or negative, respectively. This strategy leads to longer decisions when more memories have been retrieved (top). Its decision variable, (the *recalled offer value*), also differs from the *true offer value* due to the items and associated rewards that the model actually recalls on each decision (bottom), and can be used to more strongly predict choices. In contrast, the feature-based model instead sums up the value of rewards associated with each feature at encoding time and chooses with some noise at decision time, leading to neither of these predictions. **D)** The feature uncertainty manipulation used in experiments three and four. On half of the rounds, participants were informed either before or after encoding about which type of offers they would be given in the future. For experiment three, the screen showed either color or category offers, and for experiment four it showed a specific instance of either color or category (e.g., red, blue, object, etc.).

Importantly, this task could be solved using two different strategies, corresponding to either using episodic memory at choice time (an *episodic* strategy) or using incremental learning and selective attention at initial encoding (a *feature-based* strategy). To accomplish the former, participants could use episodic memory to compute an offer’s value ”on-the-fly” during their decisions by retrieving and summing the rewards associated with each offer-relevant item. Alternatively, they could instead attend to individual features during encoding and precompute a sum for each. While the episodic strategy offers more flexibility because episodes can be repurposed according to present demands, it comes at the expense of greater computation at choice time. Likewise, the feature-based strategy may yield more efficient decisions because it removes the need to reference individual episodes during decision making, but sacrifices flexibility. This is one instance of a broader tradeoff between computational efficiency and flexibility seen across learning and decision-making systems in the brain ^28,29,11,8^.

We examined the extent to which participants engaged in these two strategies using several variants of this task across five different experiments (total n = 535), finding evidence that humans use episodic memory to flexibly access features of past experience during decision making specifically when future task demands are unknown.

## 2 Results

### 2.1 Episodic memory allows access to details from past events for decision making

In experiment one, we first aimed to test whether people primarily rely on episodic memory rather than feature-based selective attention when making decisions in environments with multiple features. In this experiment, participants completed five rounds where each round consisted of four phases (**Figure 1A-B**). Participants first encoded six individual episodes, where each episode consisted of an item and its associated reward. This encoding phase was then followed by a brief 2-back working memory task to prevent active rehearsal. Immediately following this task, participants then made value-based decisions about features of earlier encoded episodes. At the end of each round, participants were asked to first freely recall all items they had seen in that round and to then report each item’s associated reward.

We hypothesized that the computational demands of tracking multiple features simultaneously would make a feature-based strategy impractical, leading participants to instead use episodic memory to compute offer values at the time of choice. We tested this hypothesis in two independent samples to ensure the replicability of our findings. Before addressing our primary question, we first examined whether participants learned to make effective decisions in the task. At the group-level, participants in both samples tended to take positive and to leave negative offers (Sample A: *M_accuracy_* = 63.6% ± 1.5, β_0_ = 0.46, 95% *HDI* = [0.33, 0.59]; Sample B: *M_accuracy_* = 60.8% ± 2.3, β_0_ = 0.36, 95% *HDI* = [0.17, 0.55]), despite substantial inter-individual variability (**Figure 2A**). Participants’ choices therefore reflected their ability to compute the value of each offer by summing over individual experiences.

**Figure 2:**
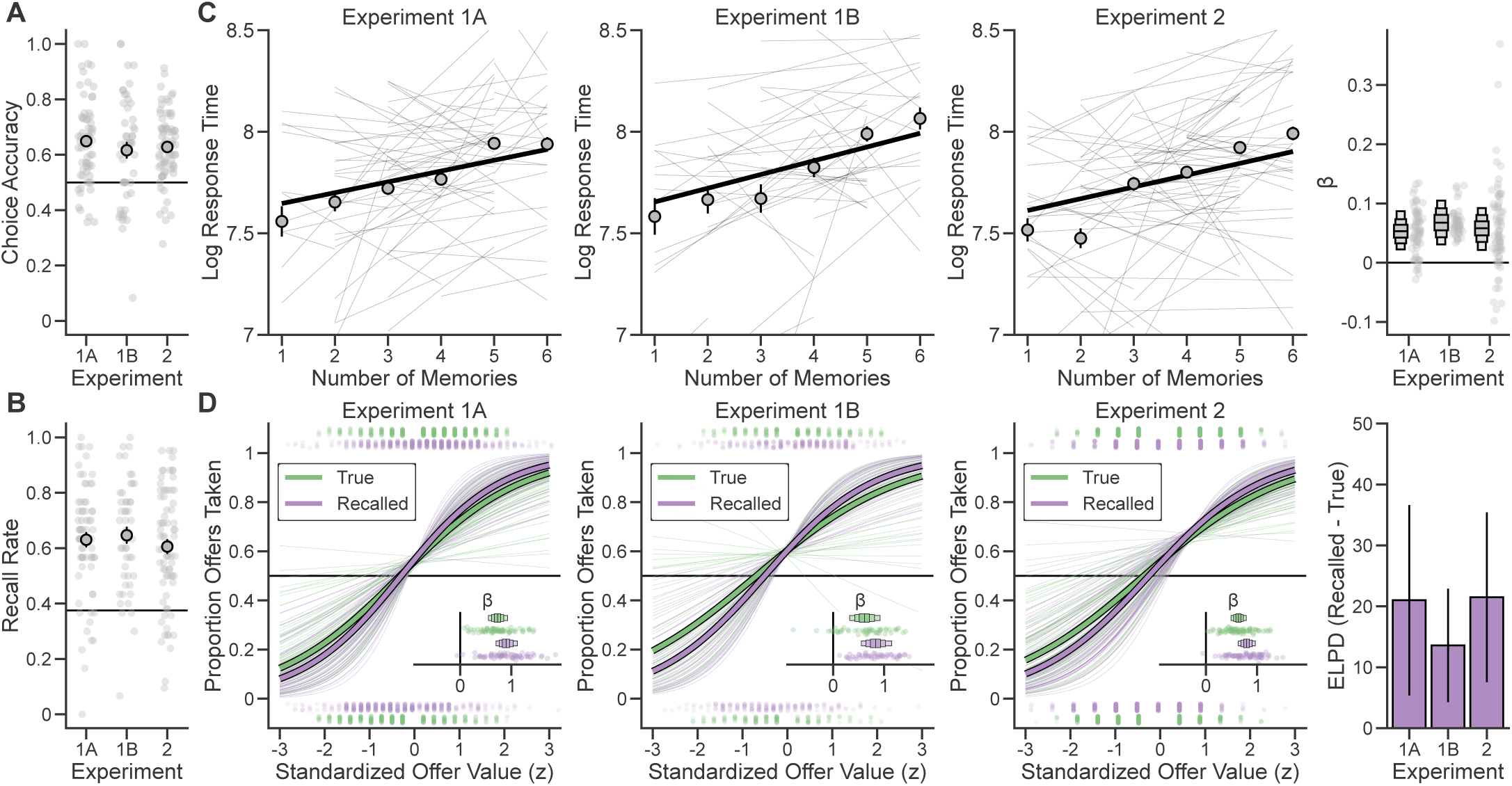
People use episodic memory to make decisions in multi-feature environments. **A)** Participants’ overall choice accuracy during the decision phase, shown for Experiment 1 (Samples A and B) and Experiment 2. Chance performance is represented by the horizontal line. Large points represent-group-level averages with error bars representing standard error. Small points represent average choice accuracy for individual participants. **B)** Participants’ rate of accurately recalling items seen in each round during the free recall portion of the memory phase. Recall rates were defined as the proportion of items that were accurately recalled relative to all items that were shown. Chance performance is represented by the horizontal line. Large points represent group-level averages with error bars representing standard error. Small points represent average recall rates for individual participants. **C)** Participants’ decision response times (log transformed) as a function of the number of memories that they accurately recalled on each round. Large lines represent group-level fits of a mixed-effects linear regression model, with fits to individuals plotted as smaller lines. Points represent group-level averages with uncertainty represented as standard error. Far right panel: Fixed effects and random effects slopes (beta) for regression model fits. Boxes represent the fixed effects posterior distribution, with horizontal lines representing the mean and boxes representing 50%, 80%, and 95% HDIs. Points represented the random effect slopes for each subject. **D)** Left: The proportion of offers that were taken as a function of summed true offer value and recalled offer value. Offer values are z-scored to facilitate comparison. Inlays show the mixed effects slopes for each predictor. Far right panel: Results of comparing the true and recalled offer value models. Model fit was first assessed using 10-fold cross-validation. The expected log pointwise predictive density (ELPD) was then computed. Higher ELPD values indicate a higher likelihood of accurately predicting new data. Here the difference in ELPD between models is shown such that positive values provide more support for the recalled offer value model. Uncertainty in the comparison is computed as in Vehtari et al. (2017) ^30^.

Our next goal was to examine whether participants maintained memory traces for the individual episodes they encoded in the first part of each round. To accomplish this, our experiment required participants to complete an additional memory phase immediately after the decision making phase of each round (**Figure 1A**). This memory phase consisted of two parts in which participants were asked to remember both elements of the episodes they had seen: they were first asked to freely recall each of the six items from a round, and then to recall the reward that was associated with each item. Participants had robust memory for the individual episodes, remembering between three and four items, on average (Sample A: *M_recall_* = 63% ± 2.5; Sample B: *M_recall_* = 64.7% ± 2.8; **Figure 2B**), which was well above chance-level recall (Sample A: β_0_ = 0.26, 95% *HDI* = [0.20, 0.31]; Sample B: β_0_ = 0.27, 95% *HDI* = [0.22, 0.33]). Item recalls exhibited classic properties of episodic memory, with items presented close together during encoding being more likely to be recalled consecutively (temporal contiguity effect ^31^; **Figure S2**), providing further evidence that participants engaged episodic memory during the task. Participants also accurately remembered the rewards associated with each item, showing a strong positive relationship between remembered and actual rewards (Sample A: β*_reward_* = 0.52, 95% *HDI* = [0.44, 0.59]; Sample B: β*_reward_* = 0.47, 95% *HDI* = [0.37, 0.56]; **Figure S3**). These results demonstrate that participants formed and retained strong memories for each episode beyond the decision making phase, and that individual memories were available for potential recall at choice time.

Next, in order to disambiguate between the episodic and feature-based strategies, we used participants’ responses on the memory phase to analyze their choice behavior. First, we reasoned that recalling an individual episode should take time ^32,33,31^ and that, accordingly, the amount of time it takes to make a decision should scale with the number of episodes that are referenced (**Figure 1C**). Importantly, the feature-based strategy makes no such prediction, as only a single item (a precomputed offer value) must be retrieved at choice time. To test this idea, we first used participants’ free recall data to determine the total number of items that they accurately recalled on each round of the task. We then examined whether they took longer to respond to offers during rounds on which they recalled more items overall. As predicted, participants took longer to decide when they subsequently recalled more memories (Sample A: β*_nMemories_* = 0.05, 95% *HDI* = [0.02, 0.08]; Sample B: β*_nMemories_* = 0.07, 95% *HDI* = [0.03, 0.11]; **Figure 2C**). We next conducted a complementary analysis examining whether decision response times were specifically related to the number of offer-relevant memories recalled — that is, only those memories whose features matched each offer (e.g., only recalled red items for offers about red things). Interestingly, we found no consistent relationship between response times and the number of offer-relevant memories recalled (see **Table S1** for results across all experiments). Together, these results suggest that participants did not retrieve exclusively the memories needed for each decision. Rather, participants broadly retrieved their memories, presumably selecting from them only the trial-relevant information before making a choice.

Having observed response time patterns suggesting that participants accessed individual memories during their decisions, we moved to examine the actual choices they made. We reasoned that if their decisions were based on the rewards they remembered being associated with each item, as predicted by an episodic strategy, their choices should be sensitive to the summed value of these remembered rewards (**Figure 1C**). To test this, we used participants’ responses on the reward memory portion of the memory phase to determine the *recalled value* of each offer. Specifically, we summed over the reported remembered reward of each offer-relevant item that was also recalled during the free recall phase. In contrast, we expected that evidence for a feature-based strategy would manifest as choices being primarily driven by the *true value* of each offer, independent of what participants explicitly remembered. This prediction follows from the nature of feature-based learning: if participants precomputed feature values during encoding, these cached values would be immune to later forgetting or distortions of individual episodes. To arbitrate between these possibilities, we fit two logistic regression models to participants’ choices: one that used the true offer value to predict each choice, and another that used recalled offer value to predict each choice. We then compared the out-of-sample predictive accuracy of each model using cross-validation.

While participants’ choices were sensitive to both true offer value (Sample A: β*_true_* = 0.73, 95% *HDI* = [0.56, 0.91]; Sample B: β*_true_* = 0.61, 95% *HDI* = [0.32, 0.92]) and recalled offer value (Sample A: β*_recalled_* = 0.89, 95% *HDI* = [0.69, 1.11]; Sample B: β*_recalled_* = 0.83, 95% *HDI* = [0.55, 1.15]), recalled offer value was slightly more effective at predicting held-out choices in both samples (Sample A: *ELPD_recalled_*_−_*_true_* = 21.00 ± 15.63, Sample B: *ELPD_recalled_*_−_*_true_* = 13.58 ± 9.30; **Figure 2D**, **Table S2**). This result indicates that participants tended to rely more heavily on the information contained in individual episodes, namely the identity and value of items, to make their decisions.

One limitation of this experiment is that it does not distinguish between two fundamentally different ways that information could be represented in memory. One possibility, which we have suggested so far, is that experiences are stored as integrated episodes, where multiple features are bound together into a single conjunctive representation (e.g., ”butterfly”) ^34,35^. Alternatively, individual experiences could be stored as separate features (e.g., ”blue” and ”animal” as independent elements) ^36^. Although both approaches could support value computation in our task, the latter becomes increasingly challenging as more features must be maintained and retrieved. To test this idea, we conducted a second experiment that was nearly identical to experiment one, but where we doubled the number of features associated with each stimulus (see **Table 3** for a summary of differences between these experiments and **Figure S1** for all stimuli used in this experiment). Specifically, in this experiment the stimuli varied across four binary features: texture (solid or pattern), location (land or sea), animacy (animal or object), and size (small or large). Offers then consisted of one of the feature levels (e.g. solid or sea) where the value of each offer was the sum of all stimuli that had the offered feature level in common (e.g. all solid things).

We reasoned that if participants relied upon memories comprised of separate features, this increased complexity would impair performance. Conversely, if participants encoded integrated episodes, the natural binding of features should preserve performance despite the additional complexity. As predicted, performance on experiment two was comparable to experiment one and all results replicated (**Figure 2**, **Table S3**), supporting the conclusion that participants relied on integrated episodic memories rather than separate feature memories to complete the task. To formally assess replicability, we pooled data across experiments one and two and tested for sample-specific deviations from the overall effects. No deviations from the pooled effects were observed across any sample (**Table S4**), providing strong evidence for the reliability of these effects.

Together, these results demonstrate that when faced with choices that could be based on many different features, participants primarily employed an episodic strategy that involved retrieving and computing over individual memories at choice time. This finding suggests that despite the increased computational demands during decision making, people prefer to maintain detailed episodic memories that can be flexibly accessed rather than attempting to track and update values for multiple features simultaneously. This preference may reflect the difficulty of implementing a feature-based strategy when faced with the curse of dimensionality, as suggested by previous theoretical work ^8^.

### 2.2 Episodic memory enables flexible decision making when it is unclear which details are important

Our findings so far demonstrate that people rely on episodic memory to make decisions in multi-feature environments, in part because episodic memories provide a natural solution to the curse of dimensionality. However, this observation alone does not fully explain why episodic memory might be specifically advantageous for decision making. To address this, we next hypothesized that episodic memory’s key benefit lies in its ability to enable flexible decisions when future task demands are uncertain — a common situation in the real world. This hypothesis predicts that people should shift away from using episodic memory when they can anticipate which features will be relevant for their upcoming decisions, as this foreknowledge would make a feature-based strategy more viable. To test this prediction directly, we next conducted an experiment where we manipulated whether participants knew in advance which features would be relevant for their decisions.

In experiment three, we contrasted two conditions: one where participants learned which features would be relevant only at choice time (*after* encoding), and another where participants knew *before* encoding which features would be needed for their upcoming decisions (**Figure 1C**). Specifically, in this new before condition, participants were told prior to encoding that they would later receive offers about color or category, but not both. This manipulation created conditions where the feature-based strategy is more feasible, as participants could safely ignore irrelevant features (e.g., category when only color offers would be made). This advantage was absent in the after condition, where feature relevance remained uncertain during encoding. We predicted that people would primarily rely on an episodic strategy in the after condition, but would shift toward a feature-based strategy when feature relevance was known in advance. We again tested this prediction across two independent samples to ensure the replicability of our findings.

Participants responded accurately, primarily taking positive and leaving negative offers both during rounds in which feature relevance was communicated before encoding (Sample A: *M* = 68.5% ± 2.9, β_0_ = 0.86, 95% *HDI* = [0.62, 1.33]; Sample B: *M* = 71.7% ± 1.9, β_0_ = 1.00, 95% *HDI* = [0.81, 1.21]) and after encoding (Sample A: *M* = 59.7% ± 2.9, β_0_ = 0.43, 95% *HDI* = [0.20, 0.66]; Sample B: *M* = 63.3% ± 2.2, β_0_ = 0.62, 95% *HDI* = [0.41, 0.83]; **Figure 3A**). Notably, participants chose more accurately in the before condition (by an average of 8.8% and 8.4% in each sample, respectively), suggesting that there were clear benefits to performance when feature relevance was known during encoding. Importantly, though, performance in the after condition was on par with our prior experiments.

**Figure 3:**
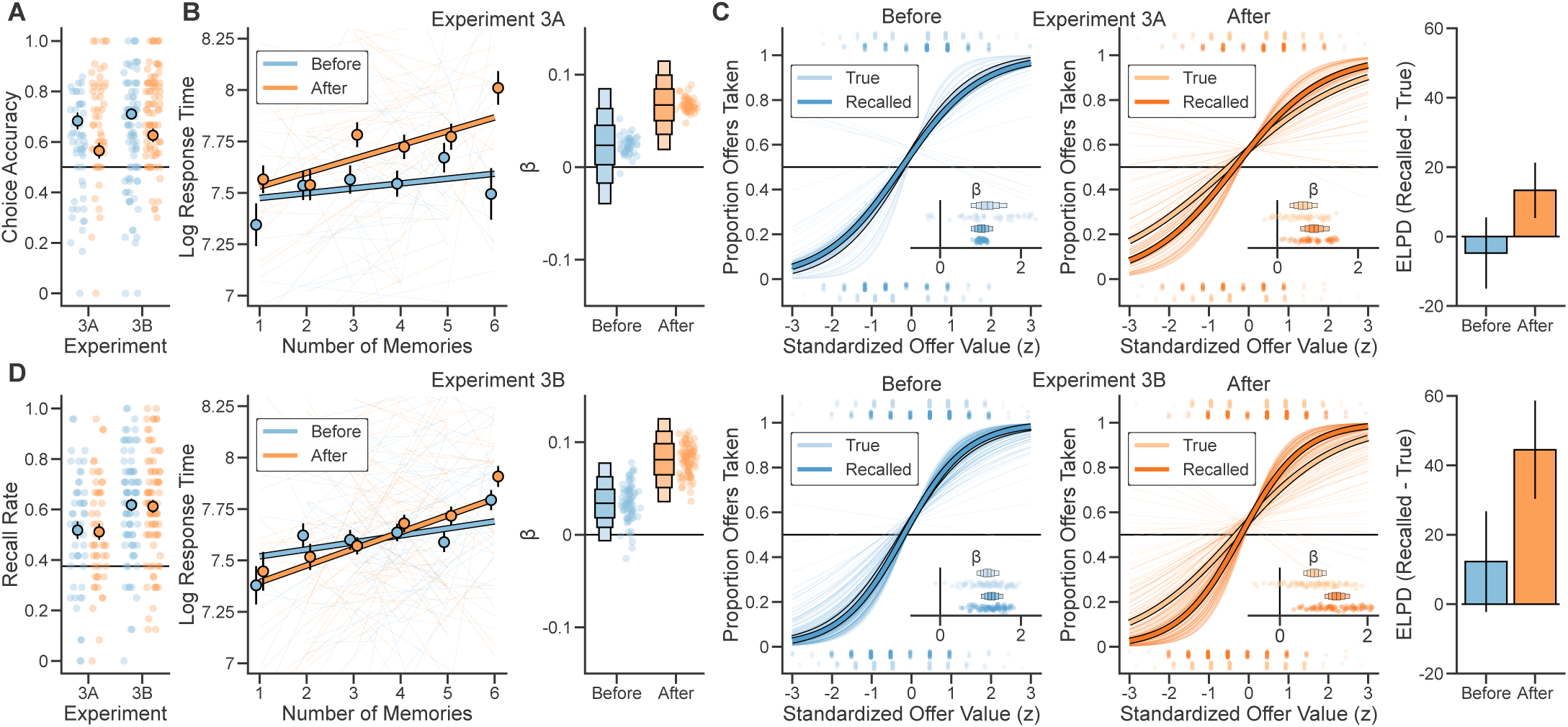
Episodic memory is primarily used for decisions when it is unclear what features are important. **A)** Participants’ choice accuracy during the decision phase, shown for Experiment 3 (samples A and B). Decisions made when the future relevance of features was known at encoding (the before condition, in blue) are shown separately from those made when this information was unknown at encoding (the after condition, in orange). Large points represent-group-level averages with error bars representing standard error. Small points represent average choice accuracy for individual participants. **B)** Participants’ decision response times (log transformed) as a function of the number of memories they accurately recalled on each round for sample A (top) and sample B (bottom). Left: Large lines represent the group-level fit of a mixed-effects regression, with fits to individuals plotted as smaller lines. Points represent group-level averages with standard error. Right: Fixed effect slopes and random effect slopes (beta) for regression model fits. Boxes represent the fixed effects posterior distribution, with horizontal lines representing the mean and boxes representing 50%, 80%, and 95% HDIs. Points represent random effect slopes for each subject. **C)** The relationship between choices and offer value for sample A (top) and sample B (bottom). Left: The proportion of offers taken as a function of summed true offer value and recalled offer value for the Before and After conditions. Mixed effects slopes are plotted as inlays. Right: Model comparison showing the difference in ELPD alongside standard error. **D)** Participants’ rate of accurately recalling items seen in each round during the free recall portion of the memory phase. Chance performance is represented by the horizontal line. Large points represent group-level averages with error bars representing standard error. Small points represent average recall rates for individual participants.

We next aimed to test our primary hypothesis. As predicted, in the after condition participants took longer to make decisions when they recalled more memories (Sample A: β*_nMemories_* = 0.08, 95% *HDI* = [0.04, 0.12]; Sample B: β*_nMemories_* = 0.06, 95% *HDI* = [0.02, 0.11]; **Figure 3B**). Yet, we found little evidence for this relationship in the before condition, when participants were aware of which features they would need for their future decisions (Sample A: β*_nMemories_* = 0.02, 95% *HDI* = [−0.04, 0.07]; Sample B: β*_nMemories_* = 0.03, 95% *HDI* = [−0.01, 0.07]). We then further examined the extent to which true and recalled offer value predicted participants’ choices in both conditions. We expected recalled offer value to be a better predictor of choices in the after condition, but not in the before condition, if participants retrieved episodes during decisions made in the former but not the latter. In line with this prediction, recalled offer value better predicted held-out choices in the after condition but not in the before condition in both samples (**Figure 3C**, **Table 1**; see **Table S2** for performance of all models). In models pooling across samples, we further assessed the interaction between these conditions, finding evidence for a greater effect of recalled memories on decision times in the After condition relative to the Before condition (β*_nMemories_*_×_*_Condition_* = 0.07, 95% *HDI* = [0.04, 0.10]; **Table S5**). Consistent with this finding, recalled offer value also provided comparably better predictions of held-out choices (β*_ELPDrecalled_*_−_*_true_*×*Condition* = 0.06, 95% *HDI* = [0.03, 0.10]). These results indicate that participants primarily referenced individual episodes during decision making when it was unclear at encoding which features would be needed for future decisions.

**Table 1:** Experiment 3 model comparison results showing fixed effects estimates (β) with 95% highest density intervals (HDI) and difference in expected log pointwise predictive density (ELPD) between true and recalled offer value models.

| Condition | Sample | Model | $\beta$ [95% HDI] | ELPD<sub>Recalled-True</sub> ± SE |
| --- | --- | --- | --- | --- |
| Before | A | True | 1.17 [0.76, 1.65] | $-4.74 \pm 10.22$ |
|  |  | Recalled | 1.02 [0.77, 1.31] |  |
| | B | True | 1.17 [0.92, 1.46] | $12.26 \pm 14.50$ |
|  |  | Recalled | 1.27 [1.02, 1.55] |  |
| After | A | True | 0.63 [0.31, 0.99] | $13.32 \pm 7.96$ |
|  |  | Recalled | 0.91 [0.58, 1.28] |  |
| | B | True | 0.79 [0.54, 1.06] | $44.50 \pm 14.13$ |
|  |  | Recalled | 1.29 [1.03, 1.59] |  |

To better understand the source of these differences in strategy recruitment, we next examined participants’ performance on the memory phase. Specifically, while participants shifted away from using episodic memory when feature relevance was known prior to decision making, it is unclear whether this effect emerged from changes in how experiences were initially encoded or from whether they were later accessed during decision making. We reasoned that if participants strategically modified their encoding based on known feature relevance, they should show impaired subsequent memory for individual episodes in the before condition, as they would be focused primarily on computing and maintaining feature-level values rather than encoding complete episodic memories. However, another possibility is that participants instead continued to encode episodic memories alongside precomputing feature values, consistent with evidence that these distinct systems often operate in parallel during learning ^25,37,38,39,40^. In this case, we expected to see comparable memory performance between conditions, with task demands influencing whether these memories were later accessed at choice time rather than whether they were initially stored.

Across both samples, we found that participants maintained strong memories of the episodes encountered in each round regardless of condition. Participants accurately recalled individual items with no differences in recall rates between conditions (Sample A: β*_a fter_*_−_*_be f ore_* = −0.01, 95% *HDI* = [−0.10, 0.08]; Sample B: β*_a fter_*_−_*_be f ore_* = −0.01, 95% *HDI* = [−0.07, 0.06]; **Figure 3D**; **Table 2**), and they further showed equally similar memory for the rewards associated with each item (Sample A: β*_a fter_*_−_*_be f ore_* = −0.06, 95% *HDI* = [−0.18, 0.06]; Sample B: β*_a fter_*_−_*_be f ore_* = −0.05, 95% *HDI* = [−0.15, 0.05]; **Figure S3**; **Table 2**). These findings demonstrate that participants encoded complete episodic memories regardless of whether they knew which features would be relevant for future decisions.

**Table 2:** Memory performance across experiments 3 and 4. Recall Rate shows mean percentage of items recalled (± standard error). β_0_ represents the intercept of a model predicting recall performance relative to chance. β*_reward_* represents the slope of a model predicting the relationship between remembered and actual rewards.

| Experiment | Sample | Condition | Recall Rate ± SE | $\beta_0$ [95% HDI] | $\beta_{reward}$ [95% HDI] |
| --- | --- | --- | --- | --- | --- |
| 3 | A | Before | 51.8% ± 3.2 | 0.14 [0.08, 0.21] | 0.55 [0.48, 0.62] |
| | | After | 51.1% ± 2.9 | 0.14 [0.08, 0.19] | 0.49 [0.39, 0.57] |
| | B | Before | 61.8% ± 2.2 | 0.24 [0.20, 0.29] | 0.53 [0.46, 0.60] |
| | | After | 61.2% ± 2.3 | 0.24 [0.19, 0.28] | 0.48 [0.41, 0.55] |
| 4 | - | Before | 62.5% ± 1.9 | 0.25 [0.21, 0.29] | 0.54 [0.49, 0.59] |
| | | After | 61.9% ± 1.9 | 0.23 [0.19, 0.27] | 0.48 [0.42, 0.54] |

Together, these results provide evidence that people selectively use episodic memory for decision making when feature relevance is unknown during encoding. When participants were aware of which features would be important for decision making prior to encoding, our results suggested that they no longer retrieved individual memories at choice time. This shift toward the feature-based strategy is sensible — it reduces computational demands at decision time by allowing direct access to pre-computed feature values rather than requiring the retrieval and integration of multiple episodic memories. This approach led to improved performance on expected decisions because the episodic strategy can introduce noise (e.g. though errors in memory retrieval), especially under time constraints when only a subset of memories might be accessible. Yet we also found that participants maintained detailed episodic memories in both conditions, suggesting that the differences we observed during decision making emerged from how information was accessed at choice time rather than how it was initially encoded. We next aimed to determine whether this parallel maintenance of episodic memories provided its own adaptive benefits for decision making.

### 2.3 Episodic memory maintains access to details if they become unexpectedly relevant

The results of experiment three suggest that while participants appeared to rely on a feature-based strategy when feature relevance was known in advance, they still maintained detailed episodic memories. One prediction that follows from this observation is that participants should still be able to use their episodic memories to maintain access to features they had initially deemed irrelevant in order to inform their decisions. In contrast, if participants abandoned episodic encoding entirely, they should struggle to access information about these irrelevant features.

We designed a fourth experiment in order to test this idea. In experiment four, once participants had completed eight rounds where the manipulation introduced by experiment three was honored, we asked them to complete a final additional round in which they were shown all possible offers. Importantly, participants were shown these *unexpected* offers regardless of whether they had been told before the encoding phase of this final round that they would see only a subset of offers later on. This allowed us to examine performance when the final round was completed under the before condition for both expected and unexpected offers. By definition, all offers shown during the after condition were unexpected. We additionally modified the task to create conditions that would encourage greater reliance on the feature-based strategy. To accomplish this, we simplified the decision phase by presenting only a single offer type per round (e.g., ”red”) prior to the final round. This modification meant that participants in the before condition needed to track only one specific feature value rather than multiple instances of the same feature (e.g., just ”red” instead of all colors), making the feature-based strategy exceptionally efficient. We predicted that this change would lead participants to rely more heavily on the feature-based strategy in the before condition relative to experiment three.

We further predicted that participants should show a specific pattern of choice behavior if they maintained access to their episodic memories despite this increased commitment to the feature-based strategy. First, in response to unexpected final round offers completed under the before condition, we predicted that participants’ accuracy should drop relative to their performance on previous rounds. This is because they would no longer be able to rely on the feature-based strategy, which provided clear benefits to performance (improving accuracy by 8-9% in experiment three when feature relevance was known in advance). Second, we predicted that to make these unexpected decisions, participants would instead need to fall back on using their episodic memories, leading to accuracy levels similar to their performance in the after condition.

We first examined overall performance on rounds prior to the final round, finding that while participants responded accurately in both conditions (Before: *M* = 78.4% ± 2.0, β_0_ = 1.57, 95% *HDI* = [1.29, 1.90]; After: *M* = 55.2% ± 2.2, β_0_ = 0.52, 95% *HDI* = [0.32, 0.74]; **Figure 4A**), they performed substantially better in the before condition relative to both samples in experiment three. Participants further showed equivalent memory performance across both conditions, with no differences in recall rates (β*_a fter_*_−_*_be f ore_* = −0.02, 95% *HDI* = [−0.07, 0.04]; **Figure 4B**; **Table 2**) or in memory for associated rewards (β*_a fter_*_−_*_be f ore_* = −0.06, 95% *HDI* = [−0.14, 0.02]; **Figure S3**; **Table 2**). These results suggest that although participants were more capable at deciding in the before condition when they had to compute only the value of a single feature, they still separately maintained episodic memories.

**Figure 4:**
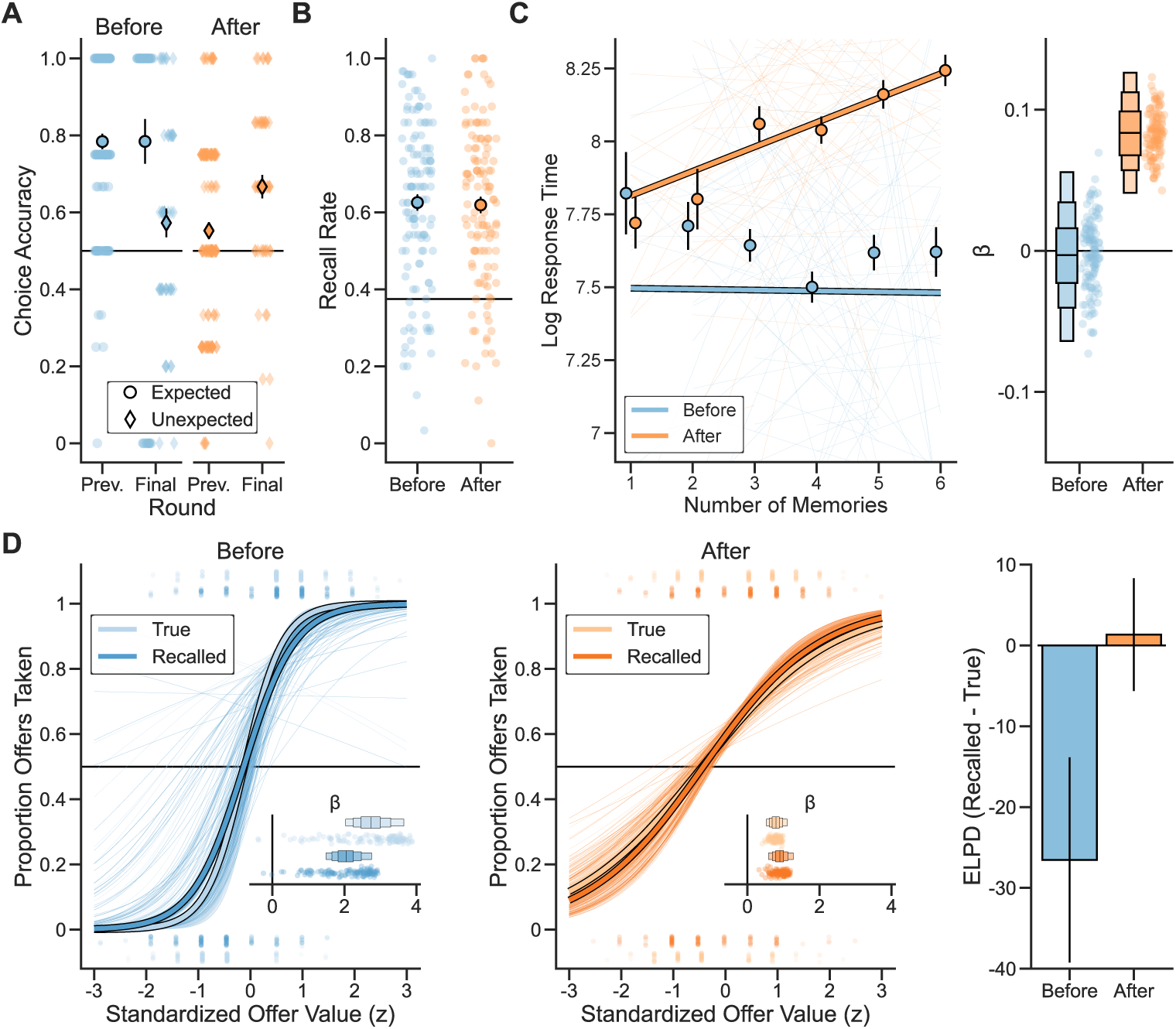
Episodic memory is used to make choices about unexpected offers. **A)** Experiment 4 choice accuracy, separated by all rounds prior to the final round (labeled as previous rounds) and the final round in which participants were asked to make decisions about both offers they expected (in the before condition) as well as those that were unexpected (in both conditions). Large points represent-group-level averages with error bars representing standard error. Small points represent average choice accuracy for individual participants. By design, only a single offer was expected in the before condition. **B)** Participants’ rate of accurately recalling items seen in each round of experiment 4 during the free recall portion of the memory phase. Chance performance is represented by the horizontal line. Large points represent group-level averages with error bars representing standard error. Small points represent average recall rates for individual participants. **C)** Experiment 4 participants’ decision response times (log transformed) as a function of the number of memories they accurately recalled on each round. Left: Large lines represent the group-level fit of a mixed-effects regression, with fits to individuals plotted as smaller lines. Points represent group-level averages with standard error. Right: Fixed effect slopes and random effect slopes for regression model fits. Boxes represent the fixed effects posterior distribution, with horizontal lines representing the mean and boxes representing 50%, 80%, and 95% HDIs. Points represent random effect slopes for each subject. **D)** Left: The proportion of offers taken as a function of summed true offer value and recalled offer value for the before and after conditions. Mixed effects slopes are plotted as inlays. Right: Model comparison showing the difference in ELPD alongside standard error.

We next asked whether this increase in performance in the before condition was due to participants’ greater reliance on the feature-based strategy. In line with this interpretation, we found that during the before condition there was virtually no evidence of a positive relationship between the number of memories recalled and decision response times (β*_nMemories_* = −0.02, 95% *HDI* = [−0.07, 0.02]; **Figure 4C**), and participants’ choices were better predicted by true offer value (β*_true_* = 2.74, 95% *HDI* = [2.03, 3.65]) than recalled offer value (β*_recalled_* = 2.04, 95% *HDI* = [1.49, 2.75], *ELPD_recalled_*_−_*_true_* = −26.54 ± 12.69; **Figure 4D**, **Table S2**). Formal comparisons confirmed that, relative to experiment three, in experiment four there was a weaker relationship between the number of recalled memories and response times in the Before condition (β_Δ_ = −0.05, 95% *HDI* = [−0.11, −0.006]), and true offer value also exerted a stronger effect on choice (β_Δ_ = 0.68, 95% *HDI* = [0.38, 0.99]). In contrast, in the after condition, participants showed longer decision response times when more memories were recalled (β*_nMemories_* = 0.09, 95% *HDI* = [0.05, 0.12]). While participants’ choices in this condition were numerically more sensitive to recalled offer value (β*_recalled_* = 0.90, 95% *HDI* = [0.59, 1.27]) than true offer value (β*_true_* = 0.79, 95% *HDI* = [0.52, 1.10]), both models were equally capable of predicting held-out choices (*ELPD_recalled_*_−_*_true_* = 1.33 ± 6.97; **Figure 4D**). This equivalent model performance likely reflects the limited data available for cross-validation, as participants made substantially fewer decisions per condition in this experiment compared to experiment three.

Finally, we turned to test the primary question of this experiment: whether in the final round participants maintained access to information about the features they were told would be irrelevant for future decisions. As predicted, participants who completed their final round in the before condition showed impaired performance on unexpected offers compared to their performance on prior rounds in this condition (*M* = 57.3% ± 3.7, β_0_ = 0.14, 95% *HDI* = [0.04, 0.24]; **Figure 4A**), but maintained high accuracy on expected offers (*M* = 78.4% ± 5.8, β_0_ = −0.07, 95% *HDI* = [−0.20, 0.06]). Critically, participants’ performance on unexpected offers matched that of the prior rounds in the after condition (β_0_ = −0.02, 95% *HDI* = [−0.11, 0.07]), suggesting they could successfully fall back on episodic memories when the feature-based strategy was insufficient. Surprisingly, we also found that participants who completed their final round in the after condition exceeded their prior performance in this condition (*M* = 66.7% ± 3.0, β_0_ = −0.13, 95% *HDI* = [−0.21, −0.05]). This improvement may stem from the increased variety of offer types shown in the final round, which provided participants with more opportunities to make decisions about items they successfully remembered, whereas previous rounds with fewer offer types may have tested only items for which their memory was weaker.

The results of experiment four suggest that even under conditions that strongly encouraged reliance on a feature-based strategy, participants maintained detailed episodic memories that they could access when needed. While participants showed clear evidence of using feature-based computations in the before condition, they were still able to rely upon their episodic memories when faced with unexpected offers, performing comparably to the after condition. This pattern of results provides evidence that episodic memory may serve as a ”backup” to aid decisions when initially irrelevant features become unexpectedly relevant.

### 2.4 Episodic memory recall becomes more targeted under realistic decision demands

An intriguing aspect of our results so far is that when participants used episodic memory for decision making, their decision times scaled with the total number of memories they recalled in each round rather than just those relevant to each offer, suggesting that retrieval during decision making was not preferentially biased toward relevant memories. Such a broad and unfocused memory search seems poorly adapted to real-world contexts where the space of possible memories is vast, raising the possibility that our relatively small memory sets may not have provided sufficient pressure for participants to develop efficient filtering strategies. In addition, our task’s simplified feature space, while allowing us to characterize episodic memory use in a controlled setting, remains far simpler than real world experience, which can consist of a vast and nearly unlimited number of features. Although participants’ choice behavior was largely unaltered by the addition of new features in experiment two, this setting was still far from naturalistic, and it remains possible that further increases to feature dimensionality could reveal more subtle differences. It is also important to note that structuring our experiments as a series of rounds – where participants knew they would be tested on their memory after each decision phase – may have encouraged them to maintain episodic memories to improve memory test performance rather than for their utility for decision making.

We designed a fifth experiment to investigate i) whether our findings would translate to a more naturalistic setting in which these constraints were relaxed, and ii) whether this would encourage participants to develop more selective retrieval strategies that improve decision making. To address these questions, we employed a larger memory set, naturalistic stimuli, and a continuous format with a surprise memory test. This task followed the structure of a single extended round in our first experiment, where participants first encoded 21 episodes that each consisted of a naturalistic image associated with a binary (+1 or -1) reward (**Figure 5A**). Following a distractor phase identical to that which was used in our previous experiments, participants were then given 30 offers about various features of the episodes (e.g. vehicles, birds, things that fly; **Figure S4**). Finally, participants completed a surprise memory test probing both free recall of the items and memory for the reward associated with each.

**Figure 5:**
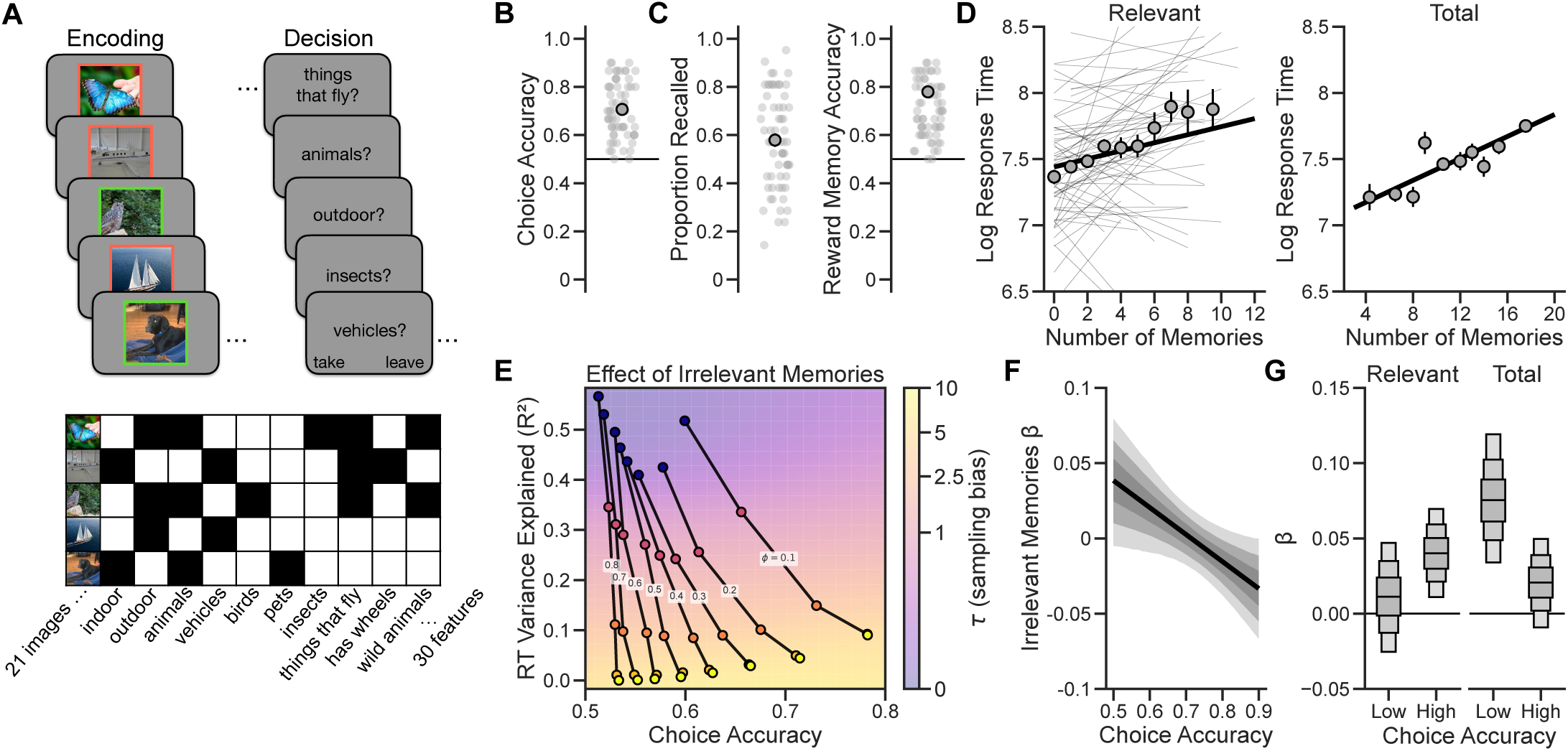
Realistic decision demands encourage participants to develop more efficient retrieval strategies. **A)** Design of Experiment 5. Top: Participants completed a single, long round of each of the four phases used in prior experiments. During encoding, participants were shown 21 naturalistic images with a binary reward (+1 or -1) represented by a green or red border. After the distractor phase, they were asked to make decisions about 30 offers about these episodes, which were then followed by a two-part memory test for free recall and the reward associated with each item. Bottom: Example features used for the offers in Experiment 5 with the images that matched each feature denoted in black. Each offer had between one and twelve matching images. See **Figure S4** for all images and offers provided to participants. **B)** Experiment 5 choice accuracy. Large points represent group-level averages with error bars representing standard error. Small points represent average choice accuracy for individual participants. **C)** Recall performance in experiment five. Left: Participants’ proportion of the 21 possible items that were accurately recalled during the free recall phase. Right: Participants’ accuracy on the two-alternative forced choice reward recall test. Large points represent group-level averages with error bars representing standard error. Small points represent average recall rates for individual participants. **D)** Experiment 5 participants’ decision response times (log transformed) as a function of the number of offer-relevant memories (left) and overall total memories (right) that they recalled. Large lines represented the group-level fit of regression models, with fits to individual subjects plotted as smaller lines. Points represent group-level averages across 10 bins with equal trials in each and standard error. Note that, because only a single memory test was completed by each participant, the total number of memories was a between-subjects measure in this experiment. **E)** Results from the behavior of simulated agents completing Experiment 5. On each offer, agents sample individual items and their associated reward without replacement, controlled by two parameters. The sampling bias (τ) determines the randomness of this sampling such that larger values bias sampling toward offer-relevant items, and a bias of zero is fully random. The probability of stopping (φ) sets a constant probability that recall will halt after each sample and controls the overall number of memories that are sampled within the task time limit. Agents with lower τ and higher φ are less accurate, and the number of offer-irrelevant memories during each decision explains a larger proportion of variance in response times when recall is more random. See **Methods** for more details. **F)** The interaction effect of the number of offer-irrelevant memories and choice accuracy on decision response times. Participants who are less accurate overall tend to have response times that are more strongly related to their number of irrelevant memories. The dark line represents the marginal posterior mean of this effect, with 50%, 80%, and 95% HDIs. **G)** Fixed effect slopes for regression models predicting the response times of median-split low and high accuracy groups from their number of offer-relevant (left) and overall total (right) memories. The horizontal line within each box represents the posterior mean and boxes represent 50%, 80%, and 95% HDIs.

We predicted that participants would demonstrate the signatures of episodic memory retrieval during decision making that we established in our prior experiments. Given the larger memory pool, we further expected that the number of offer-relevant memories recalled by participants would emerge as a stronger predictor of response times than in previous experiments, reflecting more targeted retrieval processes.

Participants performed well on this task, achieving 70.6% ± 1.0 accuracy, which was well above chance (β_0_ = 1.02, 95% *HDI* = [0.86, 1.18]; **Figure 5B**). On average, participants recalled 12 ± 0.5 items and demonstrated excellent memory for the rewards associated with each item (*M_accuracy_* = 77.9% ± 1.4; β_0_ = 0.78, 95% *HDI* = [0.74, 0.82]; **Figure 5C**). These results show that participants successfully adapted to the increased complexity of this task, maintaining both effective decision making and memory recall.

Next, we examined participants’ response times and choice behavior to test our primary hypotheses. As predicted, participants took longer to make decisions when they recalled more memories that were relevant to each offer (β*_nMemories_* = 0.03, 95% *HDI* = [0.01, 0.05]; **Figure 5D**), suggesting that their retrievals were guided by a more selective search through memory. Turning to choice behavior, we found that participants’ decisions were sensitive to both true offer value (β*_true_* = 1.03, 95% *HDI* = [0.84, 1.24]) and recalled offer value (β*_recalled_* = 1.01, 95% *HDI* = [0.81, 1.21]), consistent with our prior experiments. However, recalled offer value provided a negligible out-of-sample predictive advantage over true offer value (*ELPD_recalled_*_−_*_true_* = 4.25 ± 2.66; **Figure S4**), likely because participants’ highly accurate memory for rewards left little room for distortions in value to influence decision making.

Interestingly, we also found that participants took longer to make choices when they recalled more memories overall (β*_nMemories_* = 0.04, 95% *HDI* = [0.02, 0.06]; **Figure 5D**), similar to our previous experiments. The persistence of this effect reflects, at least in part, the fact that participants who recalled more offer-relevant memories also tended to recall more memories in general (*r* = 0.41; β*_nMemories_* = 0.74, 95% *HDI* = [0.65, 0.82]). However, a related possibility is that individual differences in memory search strategies may drive these relationships, with some participants engaging in broad, less selective retrieval that is more influenced by offer-irrelevant memories, while others employ a more targeted search process focused primarily on retrieving offer-relevant memories.

Our next aim was to investigate this idea. We predicted that task performance should be related to the selectivity of retrieval during choice. We first explored this prediction computationally by simulating agents that varied in how selectively they sampled memories during decision making. Agents that sampled more broadly and terminated search earlier showed both poorer task performance and response times that were more strongly driven by memories irrelevant to the current offer (**Figure 5E**). Consistent with this result, we found that lower-performing participants’ response times were more strongly related to the number of offer-irrelevant memories they recalled (β*_nMemories_*_×_*_Accuracy_* = −0.02, 95% *HDI* = [−0.04, −0.002]; **Figure 5F**). This finding suggests that participants who performed worse on the task engaged in less selective memory retrieval, as their decision times were disproportionately influenced by memories that were not useful for the current choice. To examine these individual differences more closely, we further split participants into low- and high-performing groups and directly compared whether total memories or offer-relevant memories better predicted response times in each group. Low-performing participants took longer to make decisions when they had more memories overall (β*_nMemories_* = 0.08, 95% *HDI* = [0.03, 0.12]), but their response times were not reliably related to the quantity of their offer-relevant memories (β*_nMemories_* = 0.01, 95% *HDI* = [−0.03, 0.05]). As expected, high-performing participants showed the opposite pattern: they took longer to respond when they had more offer-relevant memories (β*_nMemories_* = 0.04, 95% *HDI* = [0.01, 0.07]) but their response times showed no reliable relationship with their overall pool of memories (β*_nMemories_* = 0.02, 95% *HDI* = [−0.01, 0.05]). Indeed, offer-relevant memories better predicted response times out-of-sample in high-performers (*ELPD_relevant_*_−_*_total_* = 7.55 ± 3.34), confirming that these participants likely engaged in more targeted memory search during decision making, but both memory measures were equally predictive for low-performers (*ELPD_relevant_*_−_*_total_* = −0.08 ± 2.7). These findings confirm that more selective memory retrieval is associated with better decision performance, supporting the adaptive value of targeted memory search during decision making.

Together, the results of experiment five demonstrate that when memory demands more closely approximate real-world complexity – through a larger memory pool, naturalistic stimuli, and a surprise memory test – participants continued to rely on episodic memory for flexible decision making. Yet they differed in how they accessed their memories, and the ability to engage in targeted memory retrieval rather than broadly searching through all available memories distinguished better from worse decision makers. These results provide evidence that as memory demands approach real-world complexity, efficient memory search becomes a crucial determinant of decision quality.

### 2.5 Controlling for individual differences in effort

An alternative explanation for the relationship between memory retrieval and response times that we have uncovered is that individual differences in task engagement or effort may act as a confounding third variable. Under this account, more careful responding could lead to improved performance on both the decision and memory phases of our tasks, creating a spurious association between these measures rather than reflecting genuine episodic memory use during decision making.

We conducted a series of complementary analyses across each of our experiments to rule out this possibility. First, we predicted that if individual differences in effort were driving these effects, we should observe a classic speed-accuracy tradeoff during decision making. Contrary to this prediction, we found little evidence for a consistent relationship between response speed and accuracy across experiments (**Table S6**). We next used performance during the distractor phase as a proxy for effort. We reasoned that if task engagement were the common cause underlying both slower response times and better memory performance, participants with superior distractor task performance should exhibit both patterns. To test for this possibility, we included distractor performance as a covariate in our analyses. This did not meaningfully alter the strength of the observed relationship between recalled memories and response times in any experiment (**Table S7**). Lastly, we decomposed each participant’s number of recalled memories per round into within- and between-participant components to isolate effects occurring within individuals from those driven by individual differences (note that this was not possible in experiment 5, which did not employ a round-based structure). Within-participant effects remained largely consistent with our original analyses, with minor differences in the strength of effects in Experiments 1B (β*_nMemories_* = 0.05, 95% *HDI* = [−0.002, 0.10], 90%*HDI* = [0.005, 0.09]) and 4 (After condition: β*_nMemories_* = 0.05, 95% *HDI* = [−0.005, 0.09], 90%*HDI* = [0.003, 0.09]; **Table S8**). Further, when data were pooled across comparable experiments to assess replicability, the pooled mean within-participant effect remained robust, with no substantial deviations across experiments (**Table S9**).

Together, these analyses provide strong converging evidence against an effort-based explanation for our findings, supporting the conclusion that participants used episodic memory during their decisions rather than exhibiting patterns driven by individual differences in task engagement.

## 3 Discussion

Our findings indicate that people flexibly use episodic memory to guide their choices in multi-feature environments, particularly when future task demands are uncertain. When faced with multiple decision-relevant features (Experiment 1), participants relied primarily on episodic memories to compute offer values during decision making, as evidenced by both their response times and choice patterns. This strategy persisted even as feature complexity increased, suggesting that participants stored experiences as integrated episodes rather than separate feature representations (Experiment 2). When given advance knowledge of feature relevance (Experiment 3), participants shifted toward a more computationally efficient feature-based strategy that involved precomputing values during encoding. Yet despite this shift in strategy, participants continued to encode episodic memories, a parallel operation which proved useful when knowledge about previously irrelevant information was needed for decision making (Experiment 4). Finally, in an environment designed to induce pressure for participants to more efficiently recall their memories during decision making, those who retrieved more selectively made better decisions (Experiment 5). Overall, these results demonstrate that episodic memory serves as an adaptive solution to decision making under uncertainty in complex environments, allowing us to flexibly repurpose our memories according to the demands of the present.

This work connects at least two established but largely separate literatures on memory and choice. First, a number of studies focused on decision making have explored the ways in which individual experiences may be recalled for choice ^41,42,27,43,44^. This research, typically called “decision by sampling”, proposes that decision variables may be constructed by drawing samples from memory, and explains a number of ways in which peoples’ choice behavior differs when information is learned from experience rather than instructed descriptions ^45^. Much of this work has emphasized episodic memory’s value as a store for single experiences, which is a useful property when data is sparse and summary statistics cannot be reliably constructed, such as at the beginning of learning or following changes in the environment ^7,25^. Our findings address a complementary computational advantage: episodic memory’s ability to store experiences in high fidelity across multiple features simultaneously. Second, other work has proposed that episodic memory plays a critical role in our ability to infer new information about the world by allowing the formation of new links between past experiences ^46,47,48^. An important but underappreciated part of this role is episodic memory’s ability to store multiple features of experience, because each feature provides another opportunity to relate past events with one another. Here we connect each of these ideas by suggesting that features of episodes may allow for the formation of new decision variables on-the-fly, when they are needed for a choice.

Our findings add to a substantial body of research which has found that the brain contains multiple memory systems that can operate independently yet interact continuously ^39,49,40,50,11,26^, with task demands determining which is ultimately recruited for decision making ^51,52,53^. Our results suggest that this parallel operation is adaptive — while incremental learning can efficiently track relevant statistics when future demands are known, episodic memory may serve as a crucial ”backup” system by preserving detailed records of experiences. This is evidenced by participants’ ability to successfully encode detailed episodes in experiments three and four even while engaging in the feature-based strategy, and by their ability to use these memories to support unexpected decisions when incremental estimates were poorly calibrated. These findings align with recent computational work ^9^ suggesting that episodic memory may complement incrementally learned summaries by enabling the computation of new task-relevant statistics when unexpected events occur. Further research has demonstrated that episodic encoding is enhanced for schema-incongruent or surprising events ^54,55^, which suggests that creating detailed memories is most critical when current predictions fail. However, our results indicate that some level of detailed episodic encoding may occur for most experiences, providing a baseline level of flexibility for future behavior that may be further enhanced by surprise or prediction error signals ^56,57^.

Separately, our experiments uncovered systematic differences in how memory search strategies adapt to task demands and distinguish effective from ineffective decision makers. In experiments 1-4, we found that participants’ decision times scaled with their total number of memories in each round rather than just those relevant to each offer, suggesting relatively broad retrieval from memory. However, in experiment 5, we found that better-performing participants were characterized by their ability to target retrieval toward decision-relevant memories. These findings suggest that while some intrusion of irrelevant memories during retrieval is inevitable, as assumed by most computational models of free recall ^31^, people also employ more efficient search strategies to improve decision making. Future work examining how people develop selective retrieval strategies will be important to understand what factors promote more efficient memory search during decision making.

A related next step lies in characterizing *how* the various organizational principles of episodic memory guide choice. Classic computational models of memory retrieval have formalized how memories are naturally organized and accessed according to their temporal proximity and semantic relationships ^58,59,60^. This work suggests that when we retrieve a memory, it automatically triggers the recall of other memories that, for instance, occurred nearby in time or share meaningful features. Indeed, participants in our experiments showed a clear tendency to consecutively recall items that were presented close together during encoding (**Figure S3**). Such organizational principles are likely central to memory-guided decision making, as evidenced by findings demonstrating that choices are systematically influenced by temporally proximate experiences ^22^, by the context present during both encoding ^23^ and decision making ^61,23^, and by semantic similarity between experiences ^27,62^. While our results show that people can effectively use episodic memories for flexible decision making, they do not reveal the specific mechanisms by which memories are accessed. Our behavioral measures were designed to be predicted by basic properties of episodic memory recall and so remain agnostic toward any particular model or sampling algorithm. Indeed, these findings could theoretically arise from many different memory sampling strategies — from random sampling to more structured retrieval guided by these organizational principles. While we developed our toy process model to demonstrate that basic probabilistic sampling of memories can reproduce observed behavior, it likely does not capture the sophisticated retrieval mechanisms that likely govern real-world memory-guided decisions. Applying formal memory models to value-based choice could provide a more precise account of how memories are actually accessed during decision making, while potentially revealing new principles about how memory organization shapes adaptive behavior. Doing so will require not only experimental paradigms that more closely approximate real-world decision making, where people draw upon vast numbers of feature-rich experiences across extended timescales, but also methodology to measure direct memory access during decision making itself.

Related to this point, a critical feature of our experimental design was that we collected memory measures immediately following participants’ choices rather than during decision making. We did not ask participants to directly recall items during their decisions because our aim was to assess the strategy participants relied upon without instructing them to use any strategy in particular, and we reasoned that this approach may bias them toward using their episodic memories for choice. One way to circumvent this limitation would be to record neural activity during the decision phase. For example, past approaches using magnetoencephalography (MEG) have successfully decoded both the recall of individual episodes during standard memory tasks ^63,64^ and the rapid replay of sequences of memories during decision making ^65,66,67^. One possible future direction would be to similarly attempt to decode memory access during the decision phase of our task design. In addition to providing direct evidence for the recall of individual memories during choice, taking such an approach may provide multiple insights into the ways in which value is computed from memories, for instance by testing hypotheses about the number, temporal order, and semantic relationships between recalled memories.

There are also several other limitations of our experimental approach. First, we assumed that episodes consist of perfect recollections of experienced details, when in reality these details are often substantially compressed and abstracted ^68,69,70,71^. This simplification may not capture how episodic memory actually operates in naturalistic settings, where imperfect and reconstructed details may alter its relative advantages over incremental learning. Separately, our design lacked explicit behavioral measures during encoding that may have revealed more subtle differences in strategy use, particularly because the feature-based strategy should impose greater demands during encoding while the episodic strategy should be more demanding during retrieval. Future work could address this limitation by manipulating cognitive load at each of these timepoints, or by using neuroimaging to predict subsequent recall from neural activity at encoding time. Finally, while overall performance was not altered by increasing feature complexity in experiments 2 and 5, suggesting that participants relied on integrated episodes rather than separate feature stores throughout our tasks, it remains possible that people differ in the extent to which they employ either of these representations during choice. This possibility could be addressed by future experiments that manipulate feature complexity within rather than across individuals.

In conclusion, our results demonstrate that episodic memory plays an important role in enabling flexible decision making when future task demands are unknown. By maintaining detailed representations of individual experiences, episodic memory allows us to access details from our past if they become relevant for present decisions. This flexibility comes at the cost of increased computational demands during choice, leading people to adopt more efficient strategies based on precomputing decision variables when possible. Recent work on the timing of memory-based decisions supports this view, showing that people proactively compute value from memory whenever circumstances allow them to anticipate future choice requirements ^72^. Together, these findings suggest that one reason why we maintain detailed memories of the past is to help us flexibly adapt to an uncertain future.

## 4 Methods

### 4.1 Experimental Procedure

Unless otherwise noted, all procedures were identical across experiments, and differences between experiments are summarized in **Table 3**. Participants completed a four-part task over the course of a single online session designed to measure whether people access individual episodes to make decisions based on multiple features of past experiences (**Figure 1A**). Completing all four parts (a *round*) took approximately five minutes, and participants completed five (Experiment 1), seven (Experiment 2), or eight (Experiments 3 and 4) rounds in total. In Experiment 5, participants completed a single round that took approximately 20 minutes.

**Table 3:** Summary of Experimental Differences

|  | Experiment 1 | Experiment 2 | Experiment 3 | Experiment 4 | Experiment 5 |
| --- | --- | --- | --- | --- | --- |
| **Number of rounds** | 5 | 7 | 8 | 9 | 1 |
| **Number of stimulus features** | 2: color and category | 4: texture, location, animacy, size | 2: color and category | 2: color and category | 30: unspecified |
| **Reward range** | -9 to 9 | -2 to 2 | -2 to 2 | -2 to 2 | -1 to 1 |
| **Decisions per round** | Sample A: 6<br>Sample B: 3 | 4 | Sample A: 3<br>Sample B: 3 | 1 (6 in final round) | 30 |
| **Feature uncertainty manipulation** | No | No | Yes | Yes | No |
| **Surprise choice manipulation** | No | No | No | Yes | No |

#### 4.1.1 Stimuli

In Experiments 1, 3, and 4, stimuli varied across two features: color (red, yellow, blue, or green) and category (animal, object, food, scene). In Experiment 2, stimuli instead varied across four binary features: texture (with or without a pattern), location (aquatic or not aquatic), animacy (animal or object), and size (larger or smaller than a microwave). In Experiments 1-4, a total of sixteen possible items were used, with a subset of six pseudo-randomly sampled to be used in each round. All two-feature items and an example subset (highlighted in red) are shown in **Figure 1B**. As demonstrated in that figure, the six items used in each round were selected such that there were always three items with a specific instantiation of each feature (e.g. three green items, three animals), two of another type (e.g. two blue items, two objects), and one of another type (e.g. one food, one yellow). Items could repeat across rounds and the number of repetitions per item was pseudo-randomly balanced within each session. Items repeated between 1 and 4 times in a single session depending on the Experiment. Each time an item appeared it was associated with a different reward. In Experiment 5, stimuli were naturalistic images with many possible features and 21 images were shown a single time each.

#### 4.1.2 Encoding Phase

In the first part of a round, participants completed a task designed to allow them to encode individual items and an associated reward (which we refer to throughout as an *episode*). Each item was presented on the screen for 1 second, after which its reward appeared alongside it for another 6 seconds. An item’s associated reward was a pseudo-randomly sampled integer (excluding 0) between either -9 and 9 (Experiment 1), between -2 and 2 (Experiments 2, 3, and 4), or between -1 and 1 (Experiment 5). In Experiment 5, to aid future recall the reward was presented using a colored border around the presented image (green for 1 and red for -1). In order to create a balance of future offer values in each round, the reward assignment algorithm ensured that i) an equal balance of positive and negative reward was used in each round, ii) no feature dimension (e.g. blue or object) had a summed value of exactly zero, and that iii) at least one feature dimension had a positive summed value and at least one had a negative total value. Immediately after viewing the episode, participants completed an attention check consisting of the item alongside two options, either the associated reward that was just shown or another randomly selected reward. They had 3 seconds to respond. Each episode was viewed only once, for a total of six trials per round.

#### 4.1.3 Distractor Phase

Immediately following the encoding task, participants completed a 90 second distractor task to prevent active rehearsal of the episodes. This distractor consisted of a 2-back working memory task in which participants were shown one of several letters in sequence. Participants were asked to identify whether the current letter matched the one presented two steps earlier.

#### 4.1.4 Decision Phase

Immediately following the distractor task, participants then made several decisions based on the features of each item (6 decisions: Experiment 1 (sample A); 4 decisions: Experiment 2; 3 decisions: Experiments 1 (sample B), 3; 1 decision: Experiment 4; 30 decisions: Experiment 5). Each decision consisted of an offer in which a single feature (e.g. *animal*) was displayed on the screen, and participants were asked to either take or leave this offer. Participants were informed that the value of each offer consisted of the sum of each episode that was described by the offer (e.g. the value of the *animal* offer would be the sum of the rewards associated with all animals seen during encoding), and that they should take positive offers and leave negative offers. Participants had 7.5 seconds to make each decision.

#### 4.1.5 Memory Phase

Finally, immediately after the decision task, we assessed participants’ memory for the episodes in two ways. In Experiments 1-4, participants were asked to freely recall the items that they saw in each round. They were provided with six empty text boxes and were told to enter the items in the order in which they remembered them. Participants were further told to halt their recall and move on to the next task if they could no longer remember any items. Following the free recall portion, participants were shown each item and were asked to provide their memory for the reward that was associated with each item. In Experiment 5, free recall proceeded similarly to the previous experiments except that participants were given a single large box to enter their responses. Value recall in experiment 5 consisted of a two-alternative forced choice between each image with either a red or green border, and participants were instructed to choose the option that matched what they saw earlier in the task.

#### 4.1.6 Feature Uncertainty Manipulation: Experiments 3 and 4

In order to determine whether episodic memory is used preferentially when it is unclear which features should be prioritized during encoding, we manipulated whether information was provided to participants about upcoming choices in experiment 3 and 4. In these experiments, participants were told either *before* the encoding phase (4 rounds) or *after* the distractor phase (4 rounds) that they would be shown offers of only one feature type (either color or category) during the decision phase. The order in which participants were shown each condition was counterbalanced.

#### 4.1.7 Surprise Choice Manipulation: Experiment 4

We aimed to further test whether participants had access to features that were made irrelevant by the feature uncertainty manipulation, as we predicted that one hallmark of using episodic memory for decision making would be the availability of all of the features of an event at decision time. To accomplish this, we added a final round to experiment 4 where we surprised participants by telling them that regardless of whether they had been told they would only be given offers about one feature or not, they would now need to make decisions about all possible offers.

#### 4.1.8 Task Instructions and Practice

Across Experiments 1-4, participants were told a version of the following at the beginning of the experiment, with slight variations depending on the experimental condition. For brevity we have left out aspects of the instructions related to button presses and task timing:

Today you will be playing a game where your job is to earn as many points as possible. You will first see several images and will learn how many points each image is worth. You will then be asked to make choices based on what you have learned in order to earn points. The game will unfold over several different phases, and you will complete multiple rounds of these phases. Each round is independent of the others, so what you learn in one round does not influence the other rounds at all.

In the first phase, you will be shown several images, one at a time, alongside how many points it is worth. Each image can be worth either a positive or negative number of points. After viewing an image and its value, you will be asked to identify the number of points it was worth. You will be shown two numbers on either the left or right side of the screen. One of these numbers will be the image’s value, while the other will not. You will see each image and its point value only once in a round.

You will need to use the value of each image to make choices. You will be given an offer on the screen and will be asked whether you would like to take or leave the offer. You should take an offer if you think it will allow you to earn more points, and you should leave an offer if you think it will cause you to lose points. All offers will consist of different features of the images you saw. Each offer is worth the sum of the point values of all images that match that feature.

In between the two phases you just learned about, you will complete another brief task. During this phase, you will see a sequence of letters presented one at a time. Your job is to determine if the letter on the screen matches the letter that appeared two letters before.

The last phase is a brief memory test where you will be asked to recall information about the items and point values that you learned about in the first phase.

The instructions for Experiment 5 used this same language but did not include any information about rounds or the memory test.

During the instructions, participants were provided with two practice trials for each phase except the memory phase. Participants were required to achieve 100% on a multiple choice comprehension test with ten questions about the instructions in order to proceed with the study. They were further required to repeat all instructions related to any missed questions until they answered correctly. If they missed more than three questions, they were required to repeat the entire sequence of instructions, including practice trials.

#### 4.1.9 Participants

All experiments were approved by the New York University Institutional Review Board and all participants provided informed consent prior to their participation. Participants with normal or corrected-to-normal vision were recruited from the New York University subject pool. Compensation was provided in the form of course credit. Participants were excluded if they indicated on a post-task questionnaire that they wrote any information down during the study or if they either failed to answer or provided nonsensical responses to the post-task questions. To incentivize honest responses on this questionnaire, participants were told that they would receive compensation regardless of their answers.

Experiment 1 (sample A), 83 participants were recruited and 16 were excluded, leading to a final sample of 67 participants (*M_age_* = 19.05 ± 0.15; 23 males, 42 females, 2 declined to say). For Experiment 1 (sample B), 58 participants were recruited and 8 were excluded, leading to a final sample of 50 participants (*M_age_* = 18.76 ± 0.2; 14 males, 36 females). For Experiment 2, 90 participants were recruited and 15 were excluded, leading to a final sample of 75 participants (*M_age_* = 18.89 ± 0.17; 17 males, 56 females, 2 declined to say). For Experiment 3 (sample A), 61 participants were recruited and 11 were excluded, leading to a final sample of 50 participants (*M_age_* = 18.93 ± 0.18; 13 males, 37 females). For Experiment 3 (sample B), 97 participants were recruited and 13 were excluded, leading to a final sample of 84 participants (*M_age_* = 18.81 ± 0.14; 25 males, 58 females, 1 declined to say). For Experiment 4, 139 participants were recruited and 14 were excluded, leading to a final sample of 125 (*M_age_* = 19.14 ± 0.12; 42 males, 83 females). We recruited a larger sample for Experiment 4 because each participant could complete only one of the before/after conditions during the final ”surprise” round, and we aimed to have roughly comparable numbers for each condition to our prior experiments. For Experiment 5, 72 participants were recruited and 9 were excluded, leading to a final sample of 63 participants (*M_age_* = 18.78 ± 0.16; 17 males, 46 females). We determined our sample sizes based on effects measured in an initial pilot study, which is reported in the supplementary material (**Figure S5**).

### 4.2 Model Simulations

We formalized the episodic-based decision strategy using a toy process model. This model makes decisions by sequentially sampling individual memories and their associated rewards without replacement. Memories each consist of a binary feature vector and reward. When given an offer, the model first samples a memory *i* with probability computed using a softmax function with sampling bias parameter τ. The logit is defined as:

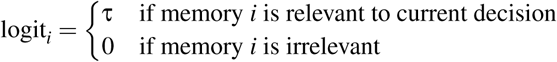

The sampling probability for each memory is then:

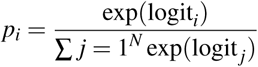

This creates a sampling bias where relevant memories have probability 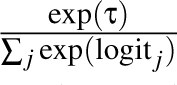 while irrelevant memories have probability 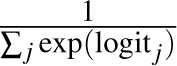. When τ = 0, all memories are sampled uniformly. As τ increases, the model increasingly favors sampling memories that *j j* share features with the current offer.

After sampling a memory, the model adds its reward to a running sum and increments the elapsed decision time by a fixed recall time. The model then decides whether to continue sampling using a geometric stopping rule: at each step, sampling terminates with probability φ. If sampling continues, the model selects another memory from the remaining unsampled memories, with probabilities renormalized over the available set. The recalled value for memory *i* includes Gaussian noise to mimic imprecise recall of reward:

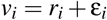

where *r_i_* is the true reward and ε*_i_* ∼ 𝒩 (0, σ^2^). The total recalled value is then:

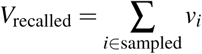

Sampling terminates when either: (1) the stopping criterion is met, (2) all memories have been sampled, or (3) maximum decision time is reached. The total decision time is then:

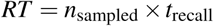

where *t*_recall_ is a constant. Finally, once sampling has terminated, the model makes a binary choice based on whether the total recalled value is positive:

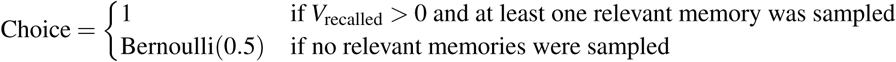

We also formalized the feature-based strategy using a model that precomputes the value of each feature during encoding by summing the rewards associated with all items sharing that feature. During the decision phase, the model simply retrieves the cached value for the offered feature and makes a choice according to a logistic choice rule with inverse temperature β. This strategy predicts constant decision times based on only on a non-retrieval time ρ since only a single cached value must be referenced on each choice.

To first illustrate the behaviors that allowed us to distinguish between episodic and feature-based decision strategies in **Figure 1C**, we simulated each model on the same basic structure used with human participants in Experiments 1-4 (*N* = 6 memories). We ran 5000 simulations for each model with the following values for each parameter. For the episodic model these were: recall time *t_recall_* = 1, reward recall noise σ = 0.5, stopping probability φ = 0.1, sampling bias τ = 0, and a maximum decision time of 7.5s. In these simulations, *t_recall_* exerts no influence over recall because its product with the number of possible sampled memories cannot exceed the maximum decision time. For the feature-based model this was: non-retrieval time ρ = 1.5s and inverse temperature β = 5.0. We chose these parameters purely to illustrate i) the expected positive relationship between decision response times and the number of sampled memories and ii) that recalled offer value should differ from true offer value as a function of the sampled memories and distortions in reward recall in the episodic memory model. In general, these properties are expected under any values of φ and σ large enough to create variation in recalls across decisions.

We also simulated episodic memory agents completing Experiment 5 (**Figure 5E**) by generating *N* = 21 memories with features chosen to create a variety of offers with different numbers of relevant memories. Performance was then simulated across several parameter combinations (sampling bias τ ∈ {0, 1, 2.5, 5, 10} and probability of stopping *p*_stop_ ∈ {0.1, 0.2, 0.3, 0.4, 0.5, 0.6, 0.7, 0.8}), with 5000 independent agents for each combination. We used an upper bound of 7.5 seconds for each decision. The reward recall noise σ was fixed to a negligible value (0.001) in order to more clearly assess the impacts of sampling bias and the probability of stopping on choice accuracy. We also fixed the recall time *t*_recall_ to 1 in order ensure that agents with minimal stopping tendencies (low φ) could not exhaustively search all available memories, thereby preserving the variance in retrievals necessary to create individual differences in performance. Because decision response times are determined by the overall number of memories recalled, we focused on assessing how each parameter combination impacts the relationship between the number of sampled memories that were irrelevant to each offer and accuracy. We calculated the amount of variance in response times explained by irrelevant memories across all of the agents with each combination of parameters.

### 4.3 Statistical Analysis

All data was analyzed with regression models estimated using hierarchical Bayesian inference such that group-level priors were used to regularize subject-level estimates unless otherwise specified. Predictors were specified as fixed effects alongside random slopes and intercepts that were allowed to vary across subjects. In experiments 3A, 3B, and 4, parameter estimates for rounds completed in either the Before or After conditions were fit separately. The joint posterior was approximated using No-U-Turn Sampling as implemented in stan ^73^. Four chains with 2000 samples (1000 discarded as burn-in) were run for a total of 4000 posterior samples per model. Chain convergence was determined by ensuring that the Gelman-Rubin statistic R was close to 1. Default weakly-informative priors implemented in the brms package were used for each regression model ^74^. For all models, fixed effects are reported in the text as the mean of each parameter’s marginal posterior distribution alongside 95% highest density intervals (HDIs), and are shown in figures as 95%, 80%, and 50% HDIs, which each indicate where that percentage of the posterior density falls. Parameter values outside of these ranges are unlikely given the model, data, and priors.

#### 4.3.1 Response Time Analysis

To examine how the number of recalled memories impacted the amount of time it took participants to make their choices in all experiments, we used a linear mixed-effects model to predict trial-wise response times. Response times were modeled using a shifted lognormal distribution, which accounts for the positive skew typical of response times. We included both random intercepts and slopes for participants to account for individual differences in both baseline response speed and sensitivity to the number of memories recalled. The model can be written as:

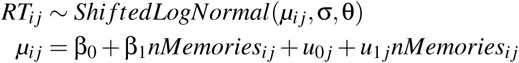

where *i* indexes trials and *j* indexes participants. The parameters *u*_0_ *_j_* and *u*_1_ *_j_* represent the random intercepts and slopes for each participant, σ is the scale parameter, and θ is the shift parameter of the distribution. The number of memories participants recalled in each round (*nMemories_i j_*) was included as a continuous predictor. We further conducted a separate analysis of the number of offer-relevant memories that participants recalled in each round, which consisted of an identical model but with this predictor instead.

In experiment 5, the mixed effects model of the total number of memories recalled by participants included only a random intercept because this measure did not vary within participants. We also assessed how individual differences in task performance related to the impact of recalled memories on response times in two ways. First, using the same mixed effects modeling framework, we modeled the relationship between the number of offer-irrelevant memories participants had and their performance as:

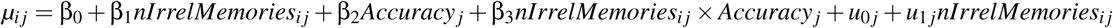

where *Accuracy_j_* is each participant’s average choice accuracy. Second, we determined whether the total number of memories or the number of offer-relevant memories better predicted response times as a function of performance. Specifically, we split participants into two groups based on the median task performance, re-fit the original mixed effects models to each group, and then assessed model fit by separating the data into 10-folds and cross validating, as described in detail in our decision analyses.

#### 4.3.2 Decision Analysis

To analyze the extent to which decision-relevant variables influence participants choices in all experiments, we fit two mixed-effects logistic regression models. Binary choices (1 = offer taken, 0 = offer rejected) were predicted by either an offer’s true value or its recalled value. The true value of an offer (*TrueValue_i j_*) was just the sum of all offer-relevant items that were shown to the participants. In contrast, we computed the recalled offer value (*RecalledValue_i j_*) as the sum over the reward that was remembered for each offer-relevant item that was also recalled during the free recall phase. These predictors were z-scored prior to model fitting in order to facilitate comparison between them. Each model included random intercepts and slopes to account for individual differences in both baseline offer acceptance rates and value sensitivity. These models can be written as:

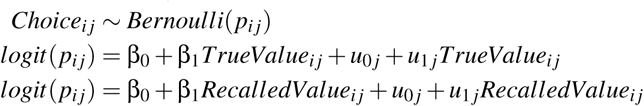

To determine the extent to which either recalled offer value or true offer value best predicted choices, we compared models fit using either predictor. Specifically, model fit was assessed by separating the data into 10-folds and cross validating. The expected log pointwise predictive density (ELPD) was then computed by summing the log likelihood for each held out datapoint and then used as a measure of out-of-sample predictive fit for each model, where higher ELPD values suggest better model fit, as they indicate a higher likelihood of accurately predicting new data. To compare models, we then subtracted the pointwise ELPD estimates and calculated the standard error of this difference to quantify uncertainty of the comparison ^30,75^.

Lastly, we assessed overall task performance using a separate mixed-effects logistic regression model. The model included only fixed and random intercepts to assess whether accuracy (1 = correct, 0 = incorrect) was from different from chance-level performance (0.5). The model was simply:

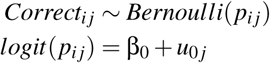

#### 4.3.3 Surprise Round Analysis

In experiment four, we also assessed performance on the final ”surprise” round where participants were required to make decisions about the offers that they expected (in the before condition) as well as for offers that were unexpected (in both conditions). To do so, we calculated a ΔPerformance score for each participant, which consisted of taking their difference in accuracy on the final round relative to their average accuracy on previous rounds (previous rounds accuracy - last round accuracy). We calculated this score separately for expected and unexpected offers in each condition, and then used ΔPerformance as the outcome variable in separate simple linear regression models:

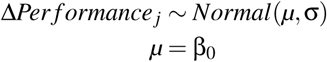

#### 4.3.4 Memory Analysis

We assessed performance on each part of the memory phase of Experiments 1-4 using two complementary models. First we examined participants’ overall recall rates, which we defined as the proportion of items they accurately recalled relative to all of the items they were shown. We compared these recall rates to chance performance, which we defined as the probability of correctly guessing items when randomly selecting 6 items from the pool of 16 possible items that could be shown on a given round, which was 37.5% (or 6/16). For each participant, we computed the difference between their mean recall rate and chance (*Recall_j_*), and then fit a simple linear regression model to these difference scores:

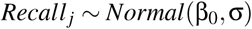

Next, to assess the accuracy of participants’ memory for the reward associated with successfully recalled items, we fit a mixed-effects linear regression model predicting participants’ remembered rewards from the true associated rewards. Both remembered (*RememberedReward_i j_*) and true (*TrueReward_i j_*) rewards were normalized by dividing by the maximum absolute value in the dataset. The model included random intercepts and slopes for participants to account for individual differences in both baseline memory and value sensitivity:

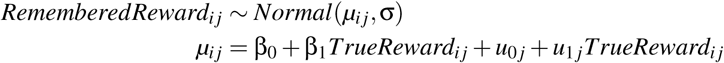

Because Experiment 5 assessed participants’ memory for associated reward using a two-alternative forced choice task, we fit a simple mixed effects model to determine the extent to which their responses differed from chance. This model was identical to the model used to assess overall task performance.

#### 4.3.5 Replication Analysis for Experiments 1-4

To assess the extent to which our effects replicated and/or differed across experiments sharing similar designs, we also ran our primary analyses with an identifier for each sample included as a fixed interaction effect. Data was pooled across comparable experiments (Experiments 1 and 2; Experiments 3 and 4), and we employed sum-to-zero (effect) coding for the experiment factor. This approach allows each experiment’s coefficient to represent its deviation from the pooled mean across all experiments, rather than a comparison to an arbitrary reference experiment. We performed these pooled analyses for both the response time and decision models. In Experiments 3 and 4, we additionally assessed interaction effects of condition on decision response times in a single mixed effects model, and assessed the difference in ELPD differences between conditions from pooled choice models. The results of these analyses are reported in **Tables S4-5**.

#### 4.3.6 Controlling for individual differences in effort

Across all experiments, we also assessed possible contributions of individual differences in effort to the relationship between the number of recalled memories and response times in three ways. First, we looked for evidence of a speed-accuracy tradeoff during the decision phase of each experiment by fitting mixed effects models predicting choice accuracy from response times:

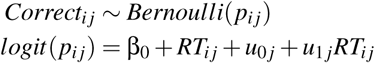

Second, we added z-scored performance on 2-back distractor task as a fixed effect covariate to the response time models. Third, to separate between-participant and within-participant effects of the number of memories shown on response times, we decomposed this predictor into two orthogonal components: a between-participant component representing each participant’s average number of memories recalled, and a within-participant component representing the deviation from each participant’s personal average. The results of these analyses are reported in **Tables S6-9**.

### 4.4 Data Availability

The data is available online at https://github.com/jonathanicholas/nm2025_emdm

### 4.5 Code Availability

The code is available online at https://github.com/jonathanicholas/nm2025_emdm

## Acknowledgments

The authors thank members of the Mattar Lab as well as Kris Jensen, Qihong Lu, Natalie Biderman, and Daphna Shohamy for their helpful comments and feedback on the project.

## 6 Supplementary Information

**Table S1:** Fixed effects estimates from regression models predicting choice response times from the number of offer-relevant memories recalled by each participant rather than the total number of memories recalled in each round, as reported in the main text.

| Experiment | Sample | Condition | $\beta$ | 50% HDI | 80% HDI | 95% HDI |
| --- | --- | --- | --- | --- | --- | --- |
| 1 | A | - | -0.07 | [-0.08, -0.06] | [-0.09, -0.05] | [-0.10, -0.04] |
|  | B | - | -0.03 | [-0.04, -0.01] | [-0.06, 0.01] | [-0.08, 0.03] |
| 2 | - | - | -0.01 | [-0.02, 0.0003] | [-0.03, 0.01] | [-0.04, 0.02] |
| 3 | A | Before | -0.07 | [-0.11, -0.03] | [-0.15, 0.002] | [-0.19, 0.04] |
|  |  | After | 0.03 | [-0.002, 0.06] | [-0.03, 0.09] | [-0.06, 0.13] |
|  | B | Before | -0.07 | [-0.09, -0.05] | [-0.11, -0.02] | [-0.14, 0.001] |
|  |  | After | -0.05 | [-0.08, -0.03] | [-0.10, -0.01] | [-0.13, 0.02] |
| 4 | - | Before | -0.10 | [-0.13, -0.07] | [-0.16, -0.04] | [-0.20, -0.005] |
|  |  | After | -0.03 | [-0.05, 0.001] | [-0.07, 0.02] | [-0.10, 0.05] |

**Table S2:** Expected log posterior density resulting from 10-fold cross validation for each choice model in all experiments.

| Experiment | Sample | Condition | Model | ELPD ± SE |
| --- | --- | --- | --- | --- |
| 1 | A | - | True | -877.00 ± 13.25 |
| | | | Recalled | -857.59 ± 14.32 |
| | B | - | True | -355.79 ± 8.35 |
| | | | Recalled | -342.21 ± 8.54 |
| 2 | - | - | True | -1062.25 ± 13.27 |
| | | | Recalled | -1040.77 ± 14.37 |
| 3 | A | Before | True | -240.82 ± 9.32 |
| | | | Recalled | -245.56 ± 8.90 |
| | | After | True | -256.41 ± 6.68 |
| | | | Recalled | -243.09 ± 7.56 |
| | B | Before | True | -446.73 ± 13.33 |
| | | | Recalled | -434.47 ± 13.31 |
| | | After | True | -485.72 ± 10.55 |
| | | | Recalled | -441.23 ± 13.01 |
| 4 | - | Before | True | -182.12 ± 13.12 |
| | | | Recalled | -208.66 ± 11.29 |
| | | After | True | -221.67 ± 6.70 |
| | | | Recalled | -220.33 ± 7.49 |

**Table S3:** Results of Experiment 2. Choice Accuracy shows mean percentage of correct choices (± standard error). Response Time shows the relationship between decision time and number of memories recalled. Choice Model Comparison shows the difference in ELPD between recalled and true offer value models. Memory measures show free recall performance and accuracy of reward memory.

| Measure | Value | 95% HDI |
| --- | --- | --- |
| Choice Accuracy | 62.4% ± 1.3 | - |
| Choice Model Intercept ($\beta_0$) | 0.45 | [0.34, 0.55] |
| Response Time Slope ($\beta_{nMemories}$) | 0.06 | [0.02, 0.09] |
| True Offer Value Slope ($\beta_{true}$) | 0.64 | [0.49, 0.79] |
| Recalled Offer Value Slope ($\beta_{recalled}$) | 0.80 | [0.63, 0.96] |
| Choice Model Comparison ($ELPD_{recalled-true}$) | 21.48 ± 13.93 | - |
| Memory Recall Rate | 60.6% ± 2.4 | - |
| Memory Recall Model Intercept ($\beta_0$) | 0.23 | [0.18, 0.28] |
| Reward Memory Slope ($\beta_{reward}$) | 0.53 | [0.47, 0.60] |

**Table S4:** Results of Replication analysis for Experiments 1-2. RT and choice show mean effects pooled across all samples and sample-specific deviations. Values show posterior means with 95% HDI in brackets.

| Sample | RT | Choice |
| --- | --- | --- |
| Pooled Mean | $\beta = 0.06$<br>95% HDI [0.05, 0.07] | $\beta_{\text{true}} = 0.64$, 95% HDI [0.53, 0.75]<br>$\beta_{\text{recalled}} = 0.82$, 95% HDI [0.70, 0.94] |
| *Deviations from Pooled Mean* |  |  |
| 1A | $\Delta\beta = -0.003$<br>95% HDI [-0.03, 0.02] | $\Delta\beta_{\text{true}} = 0.08$, 95% HDI [-0.07, 0.22]<br>$\Delta\beta_{\text{recalled}} = 0.05$, 95% HDI [-0.11, 0.20] |
| 1B | $\Delta\beta = 0.01$<br>95% HDI [-0.03, 0.04] | $\Delta\beta_{\text{true}} = -0.08$, 95% HDI [-0.24, 0.09]<br>$\Delta\beta_{\text{recalled}} = -0.02$, 95% HDI [-0.20, 0.17] |
| 2 | $\Delta\beta = -0.003$<br>95% HDI [-0.03, 0.02] | $\Delta\beta_{\text{true}} = -0.001$, 95% HDI [-0.14, 0.14]<br>$\Delta\beta_{\text{recalled}} = -0.03$, 95% HDI [-0.18, 0.13] |

**Table S5:** Results of Replication analysis for Experiments 3-4. RT and choice show mean effects pooled across all samples and sample-specific deviations. Values show posterior means with 95% HDI in brackets.

| Sample | Condition | RT | Choice |
| --- | --- | --- | --- |
| Pooled Mean | Before | $\beta = 0.007$<br>95% HDI [-0.017, 0.030] | $\beta_{\text{true}} = 1.51$, 95% HDI [1.43, 1.58]<br>$\beta_{\text{recalled}} = 1.30$, 95% HDI [1.23, 1.36] |
| | After | $\beta = 0.076$<br>95% HDI [0.053, 0.098] | $\beta_{\text{true}} = 0.75$, 95% HDI [0.69, 0.80]<br>$\beta_{\text{recalled}} = 1.03$, 95% HDI [0.96, 1.09] |
| | Interaction | $\beta = 0.069$<br>95% HDI [0.039, 0.098] | — |
| *Deviations from Pooled Mean* |  |  |  |
| 3A | Before | $\Delta\beta = 0.03$<br>95% HDI [-0.02, 0.08] | $\Delta\beta_{\text{true}} = -0.40$, 95% HDI [-0.70, -0.11]<br>$\Delta\beta_{\text{recalled}} = -0.22$, 95% HDI [-0.47, 0.03] |
| | After | $\Delta\beta = 0.01$<br>95% HDI [-0.02, 0.06] | $\Delta\beta_{\text{true}} = -0.12$, 95% HDI [-0.36, 0.13]<br>$\Delta\beta_{\text{recalled}} = -0.12$, 95% HDI [-0.38, 0.14] |
| | Interaction | $\Delta\beta = -0.01$<br>95% HDI [-0.07, 0.05] | — |
| 3B | Before | $\Delta\beta = 0.03$<br>95% HDI [-0.01, 0.06] | $\Delta\beta_{\text{true}} = -0.28$, 95% HDI [-0.55, -0.03]<br>$\Delta\beta_{\text{recalled}} = -0.04$, 95% HDI [-0.26, 0.19] |
| | After | $\Delta\beta = -0.02$<br>95% HDI [-0.07, 0.01] | $\Delta\beta_{\text{true}} = 0.01$, 95% HDI [-0.20, 0.22]<br>$\Delta\beta_{\text{recalled}} = 0.23$, 95% HDI [0.005, 0.46] |
| | Interaction | $\Delta\beta = -0.05$<br>95% HDI [-0.11, -0.0004] | — |
| 4 | Before | $\Delta\beta = -0.05$<br>95% HDI [-0.11, -0.006] | $\Delta\beta_{\text{true}} = 0.68$, 95% HDI [0.38, 0.99]<br>$\Delta\beta_{\text{recalled}} = 0.26$, 95% HDI [0.01, 0.52] |
| | After | $\Delta\beta = 0.01$<br>95% HDI [-0.03, 0.05] | $\Delta\beta_{\text{true}} = 0.11$, 95% HDI [-0.12, 0.34]<br>$\Delta\beta_{\text{recalled}} = -0.11$, 95% HDI [-0.35, 0.12] |
| | Interaction | $\Delta\beta = 0.06$<br>95% HDI [0.005, 0.13] | — |

**Table S6:** Results of regression analyses testing for the relationship of speed and accuracy in each experiment. A positive relationship was found in only the After condition of Experiment 4. Betas are the posterior mean of the fixed effects of log response time on choice accuracy, with HDIs in brackets.

| Experiment | Condition | $\beta$ | 95% HDI | 90% HDI | 80% HDI |
| --- | --- | --- | --- | --- | --- |
| 1A | — | -0.02 | [-0.18, 0.15] | [-0.16, 0.12] | [-0.12, 0.09] |
| 1B | — | -0.18 | [-0.44, 0.07] | [-0.40, 0.03] | [-0.35, -0.01] |
| 2 | — | -0.03 | [-0.19, 0.12] | [-0.16, 0.10] | [-0.13, 0.07] |
| 3A | Before | -0.31 | [-0.61, -0.03] | [-0.56, -0.08] | [-0.50, -0.13] |
|  | After | -0.04 | [-0.35, 0.26] | [-0.30, 0.21] | [-0.24, 0.15] |
| 3B | Before | -0.23 | [-0.47, 0.01] | [-0.43, -0.02] | [-0.39, -0.07] |
|  | After | -0.22 | [-0.45, 0.01] | [-0.42, -0.02] | [-0.37, -0.06] |
| 4 | Before | -1.24 | [-1.70, -0.80] | [-1.64, -0.87] | [-1.55, -0.94] |
|  | After | 0.55 | [0.13, 0.96] | [0.20, 0.89] | [0.28, 0.82] |
| 5 | — | -0.36 | [-0.48, -0.20] | [-0.53, -0.23] | [-0.42, -0.24] |

**Table S7:** Results of regression analyses predicting decision response times from the number of recalled memories alongside distractor performance as a covariate. There were no substantial changes to memory-related effects. A positive relationship between distractor performance and response times was seen in Experiments 1B and 2. Betas are the posterior mean of fixed effects, with 95% HDIs in brackets.

| Experiment | Condition | Effect | $\beta$ | 95% HDI |
| --- | --- | --- | --- | --- |
| 1A | — | # of Memories | 0.05 | [0.02, 0.08] |
|  |  | Distractor performance | 0.05 | [-0.02, 0.12] |
| 1B | — | # of Memories | 0.06 | [0.02, 0.10] |
|  |  | Distractor performance | 0.09 | [0.01, 0.18] |
| 2 | — | # of Memories | 0.06 | [0.02, 0.09] |
|  |  | Distractor performance | 0.08 | [0.01, 0.15] |
| 3A | Before | # of Memories | 0.04 | [-0.02, 0.10] |
|  |  | Distractor performance | -0.12 | [-0.23, -0.02] |
| 3A | After | # of Memories | 0.07 | [0.02, 0.12] |
|  |  | Distractor performance | -0.04 | [-0.15, 0.05] |
| 3B | Before | # of Memories | 0.04 | [-0.003, 0.08] |
|  |  | Distractor performance | -0.02 | [-0.09, 0.05] |
| 3B | After | # of Memories | 0.07 | [0.03, 0.11] |
|  |  | Distractor performance | -0.002 | [-0.07, 0.07] |
| 4 | Before | # of Memories | 0.01 | [-0.05, 0.06] |
|  |  | Distractor performance | -0.13 | [-0.21, -0.04] |
| 4 | After | # of Memories | 0.08 | [0.04, 0.12] |
|  |  | Distractor performance | 0.04 | [-0.03, 0.10] |
| 5 | — | # of Memories | 0.04 | [0.02, 0.06] |
|  |  | # of Relevant Memories | 0.03 | [0.01, 0.05] |
|  |  | Distractor performance | 0.02 | [-0.08, 0.11] |

**Table S8:** Results showing within-participant isolated effects for regressions predicting decision response times from the number of recalled memories. This predictor is the result of centering each participants number of recalled memories by their own mean. Betas are the posterior mean of fixed effects, with HDIs in brackets.

| Experiment | Condition | $\beta$ | 95% HDI | 90% HDI | 80% HDI |
| --- | --- | --- | --- | --- | --- |
| 1A | — | 0.04 | [0.004, 0.08] | [0.009, 0.07] | [0.02, 0.06] |
|  | — | 0.05 | [-0.002, 0.10] | [0.005, 0.09] | [0.01, 0.08] |
| 2 | — | 0.04 | [0.005, 0.07] | [0.01, 0.07] | [0.02, 0.06] |
| 3A | Before | 0.001 | [-0.05, 0.05] | [-0.04, 0.04] | [-0.03, 0.03] |
|  | After | 0.08 | [0.03, 0.13] | [0.04, 0.12] | [0.05, 0.11] |
| 3B | Before | 0.03 | [-0.006, 0.07] | [-0.001, 0.06] | [0.006, 0.05] |
|  | After | 0.05 | [0.006, 0.09] | [0.01, 0.08] | [0.02, 0.07] |
| 4 | Before | -0.01 | [-0.07, 0.04] | [-0.06, 0.03] | [-0.05, 0.02] |
|  | After | 0.05 | [-0.005, 0.09] | [0.003, 0.09] | [0.01, 0.08] |

**Table S9:** Results showing pooled means and sample-specific deviations of within-participant isolated effects for regressions predicting decision response times from the number of recalled memories.

| Sample | Condition | $\beta$ | 95% HDI |
| --- | --- | --- | --- |
| *Pooled Mean* |  |  |  |
| Experiments 1-2 | — | 0.04 | [0.02, 0.06] |
| Experiments 3-4 | Before | 0.01 | [-0.04, 0.05] |
|  | After | 0.06 | [0.01, 0.10] |
| *Deviations from Pooled Mean* |  |  |  |
| 1A | — | -0.002 | [-0.03, 0.03] |
| 1B | — | 0.01 | [-0.03, 0.04] |
| 2 | — | -0.004 | [-0.03, 0.03] |
| 3A | Before | -0.01 | [-0.05, 0.03] |
|  | After | 0.02 | [-0.02, 0.07] |
| 3B | Before | 0.02 | [-0.02, 0.07] |
|  | After | -0.01 | [-0.05, 0.02] |
| 4 | Before | -0.02 | [-0.07, 0.03] |
|  | After | -0.01 | [-0.05, 0.02] |

**Figure S1:**
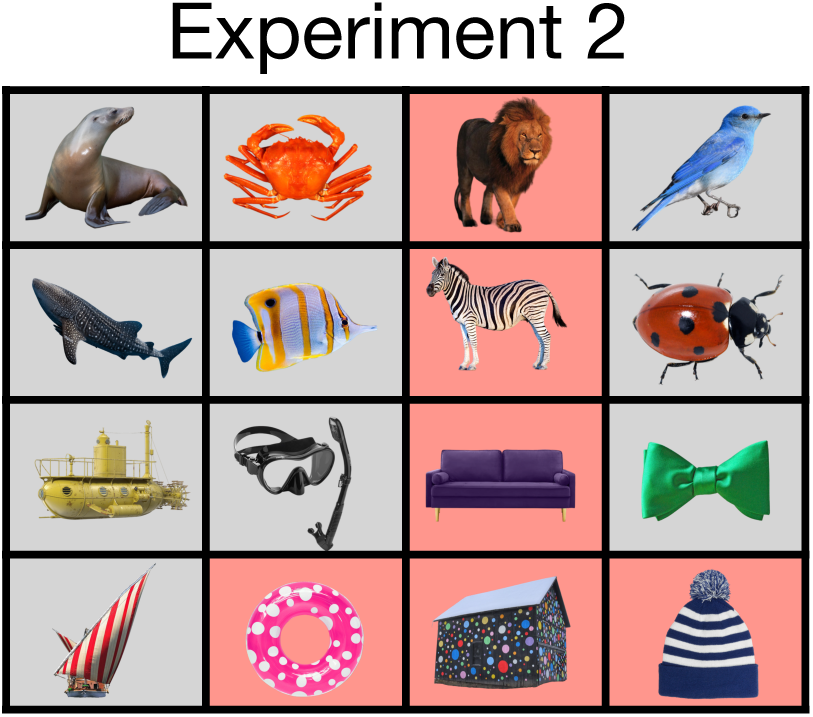
The full set of stimuli used in experiment two, which differed along four dimensions: texture (solid or patterned), location (land or sea), animacy (animal or object), and size (small or large). Six images were again sampled to be shown in each round (with an example in red).

**Figure S2:**
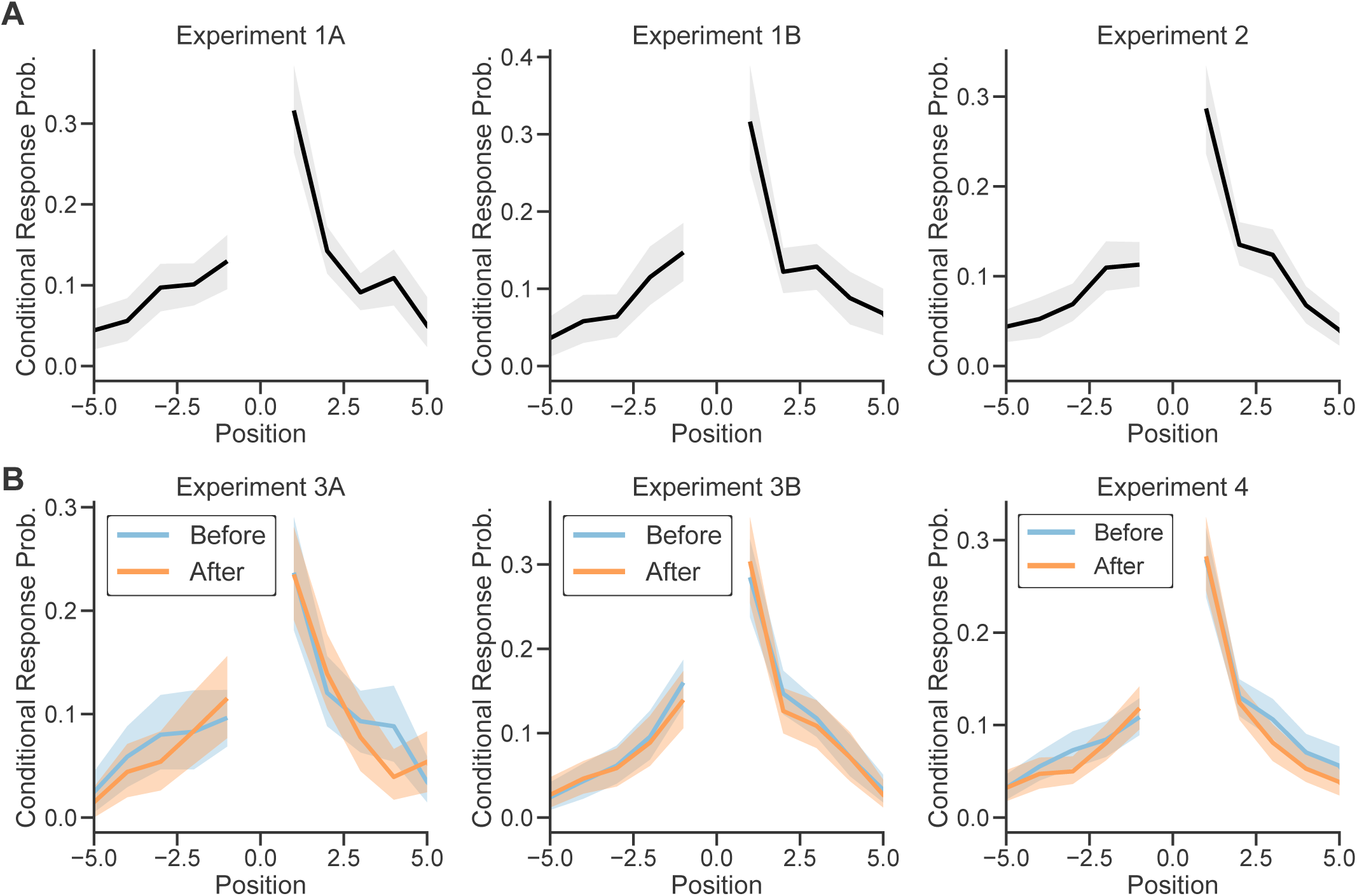
Lag-conditional response probability (CRP) curves demonstrating the classic contiguity effect (in which items encoded closer in time to one another are recalled more closely together in time ^31^) in free recall data for experiments one and two (**A**) and experiments three and four (**B**). Lines represent group-level averages and bands represent 95% CIs.

**Figure S3:**
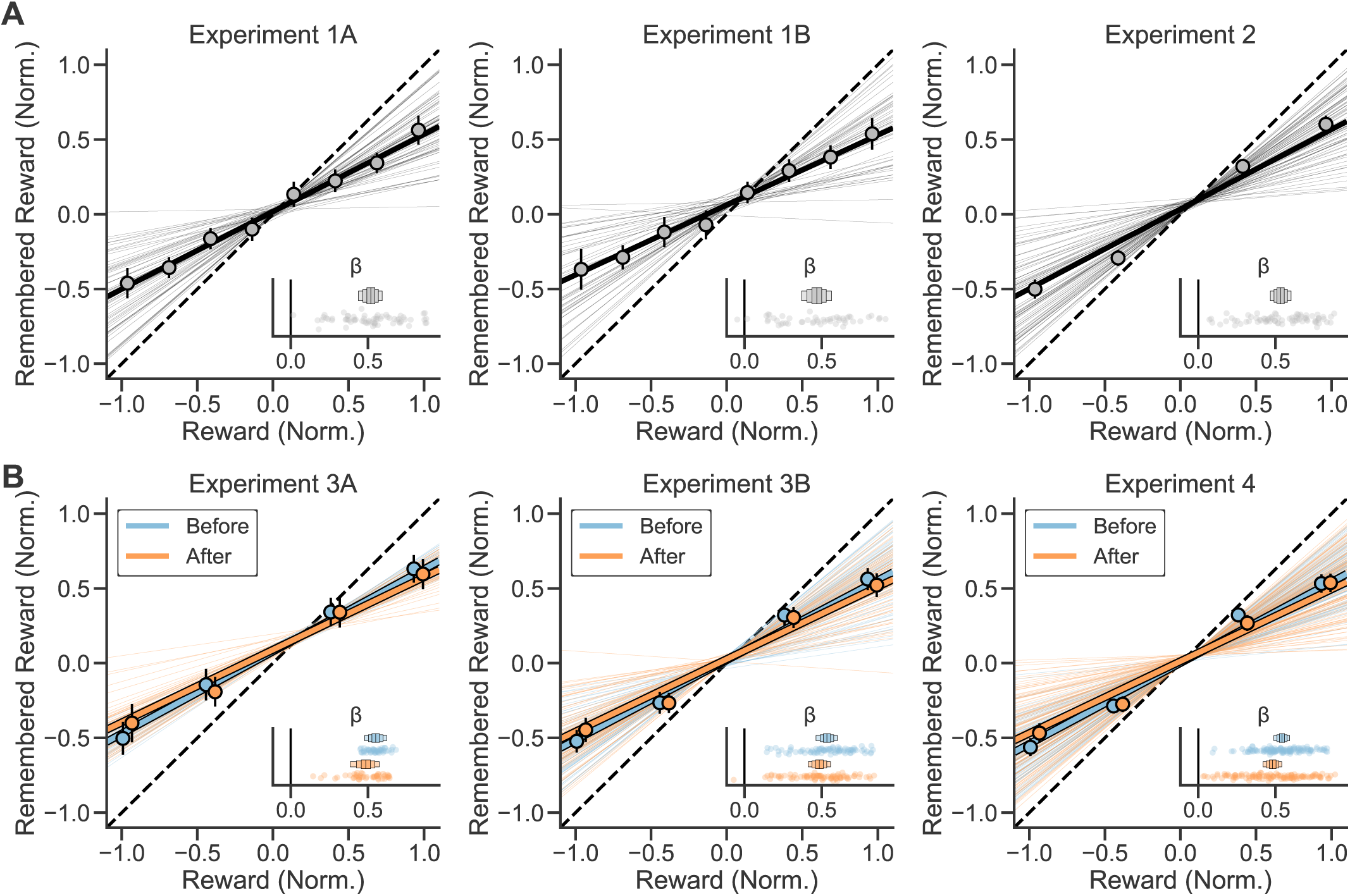
Participants’ remembered reward reported for each item during the reward memory portion of the memory phase for experiments one and two (**A**) and experiments three and four (**B**). The remembered reward for each item is plotted as a function of the item’s true associated reward. Large lines represent the group-level fit of a mixed-effects regression, with individual subject fits plotted as smaller lines. Points represent group-level averages with standard error. Inlays show fixed and random effects, where boxes represent the fixed effects posterior distribution with horizontal lines representing the mean and boxes representing 50%, 80%, and 95% HDIs. Points represent random effects slopes for each subject.

**Figure S4:**
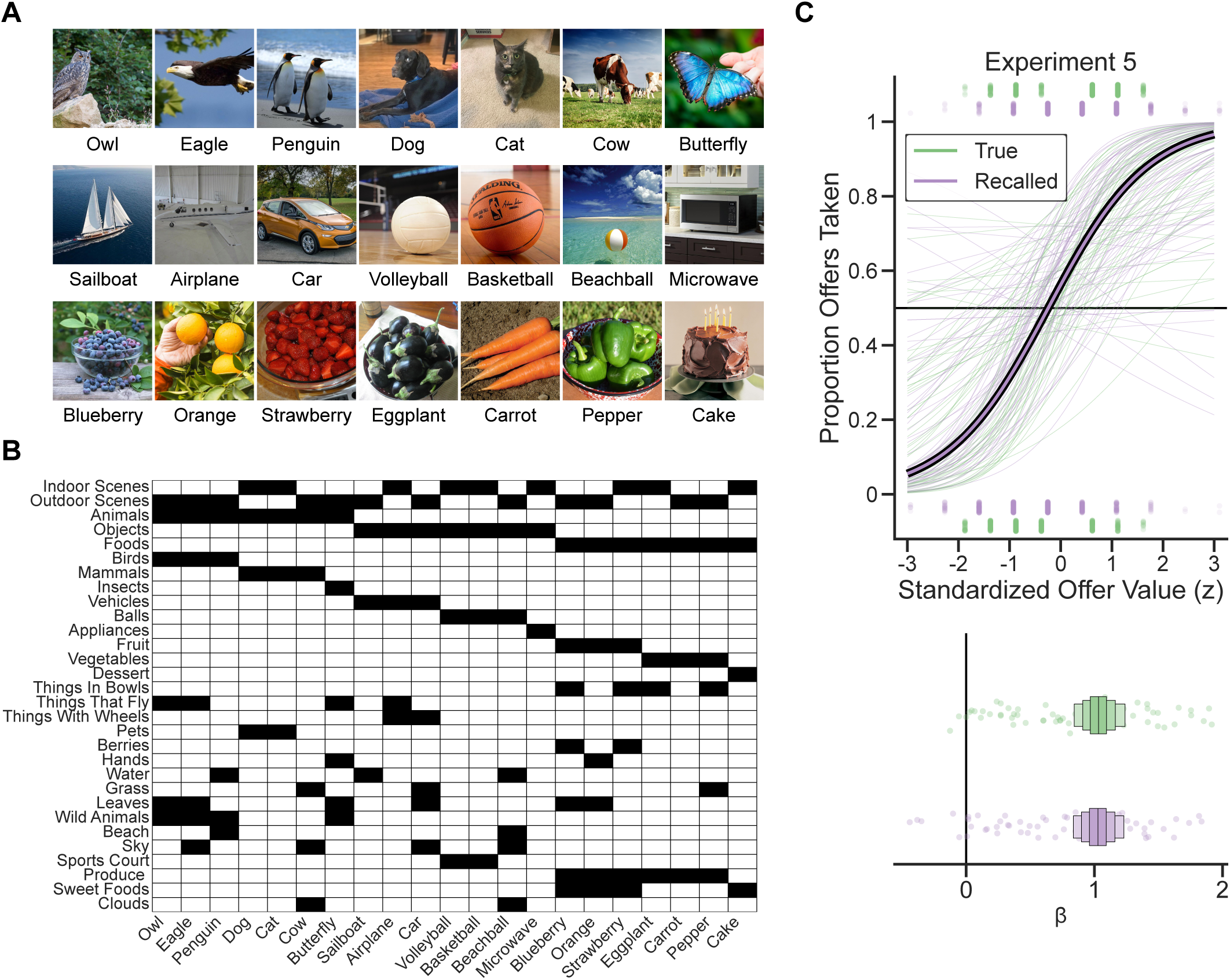
Experiment 5 Supplementary Information. **A)** The full set of images used in experiment 5. **B)** The full set of features and the images that match each feature (marked in black). **C** Top: The proportion of offers taken as a function of summed true offer value and recalled offer value for Experiment 5. Offer values are z-scored to facilitate comparison. Large lines represent group-level fits of a mixed-effects logistic regression model. Small lines represent regressions fit to individuals for visualization purposes. Bottom: Fixed and random effects for each predictor with horizontal lines represnting mean and boxes representing 50%, 80%, and 95% HDIs. True and recalled offer value were largely similar predictors in Experiment 5.

**Figure S5:**
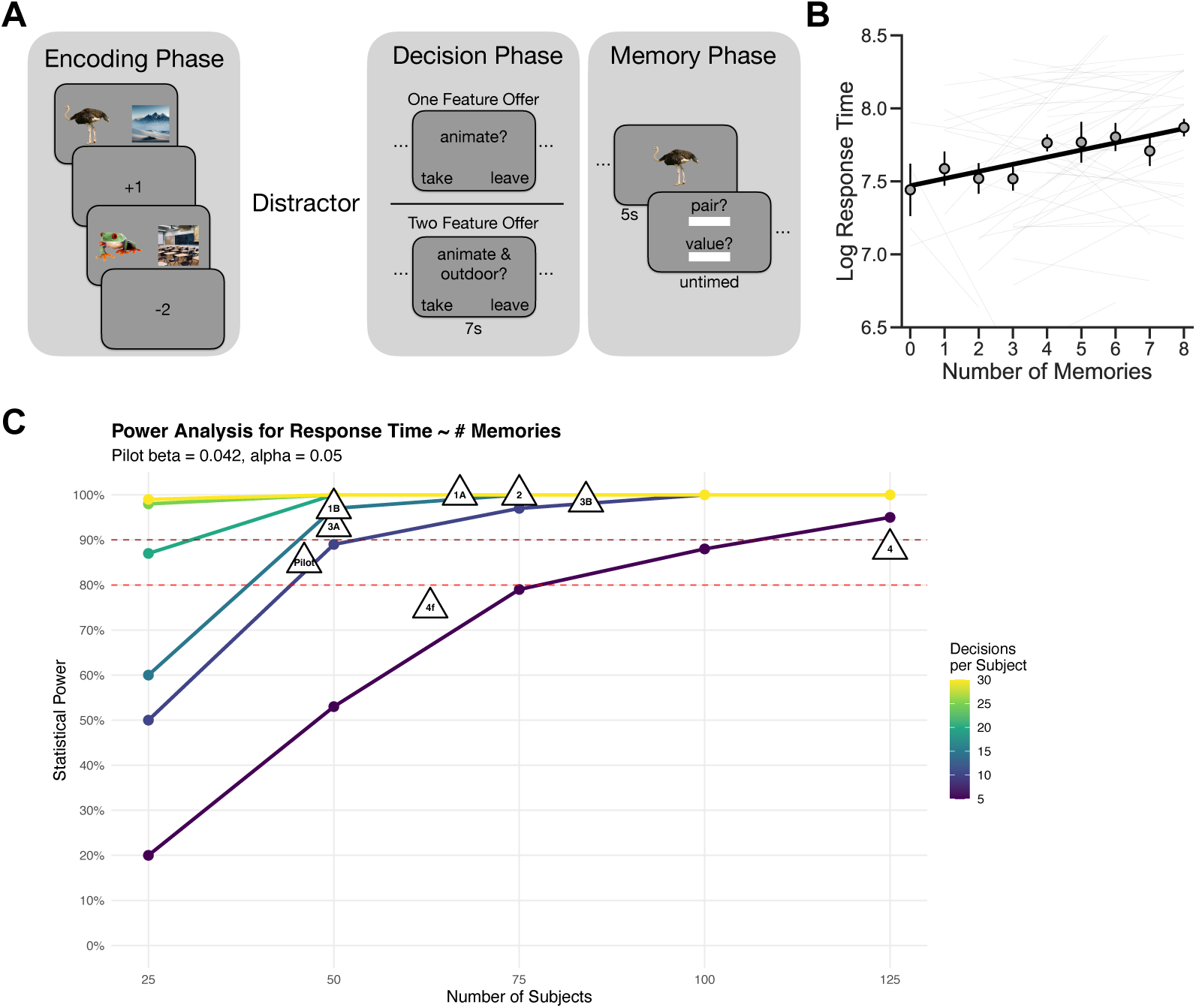
Pilot Experiment Design and Sample Size Selection. **A)** Prior to Experiment 1, we conducted a pilot study (n = 46) which was used to establish the initial effect of the number of recalled memories on decision response times and to guide future sample size selection. The pilot study consisted of four phases. During the encoding phase, participants were first shown several pairs of images alongside a value that was assigned to the pair. To encourage episodic encoding, participants were told to imagine a brief but vivid interaction occurring between the images. Each pair consisted of a scene presented alongside an object, where each scene was either an indoor or an outdoor image while each object was either an animate or an inanimate image. Each pair was shown only a single time per round. After ecnoding, participants completed a distractor that was identical that used in the experiments of the main text. Next, during the decision phase, participants were shown offers about one feature (e.g. ”indoor, ”animate”) or two features (e.g. ”indoor and ”animate”, ”outdoor and animate”). Offer values were computed the same way as in the main text experiments. Finally, during the memory phase, participants were shown one image from each pair that they encoded during the round and were asked to recall the name of the image it was paired with, alongside the value of the pair. Participants encoded eight pairs per round, and they completed four rounds of this task. In each round they made four choices. **B)** The effect of the number of recalled memories on decision response times (β = 0.04, 95%*HDI* = [0.02, 0.07]) in the pilot experiment. Note that we also predicted that single-feature decisions would take longer two feature decisions because they involve twice as many memories, but this was not supported (β = 0.13, 95%*HDI* = [−0.004, 0.26]). We speculated that this was because of differences in processing the words on the screen, but abandoned this element of the design in favor of the design used in Experiment 1. **C)** Results of a power analysis based on the effect size measured in the pilot data. The analysis was conducted by using forward simulation to model prospective studies with new participants by creating synthetic datasets with varying sample sizes and numbers of observations per participant while also preserving the varaince structure (random effects and residual variance) that was observed in our pilot data. For each scenario, we generated balanced datasets with specified numbers of participants and observations per participant, simulated a realistic range of recalled memories, and estimated statistical power using Monte Carlo simulation via the simr package. This analysis revealed that approximately 50 participants with at least 10 decisions per participant would be adequate to detect this effect with 80-90% power. We used this as a minimum bound rather than a fixed target sample size for several reasons: i) each experiment involved changes that could affect power (e.g. Experiment 2 added more stimulus features), ii) we wanted to ensure adequate power for other analyses such as choice model comparisons between true and recalled offer values, and iii) we aimed to ensure robust detection in newly added experimental conditions. We observed effects of the number of recalled memories on response times that were similar in magnitude to the pilot study across all of our experiments, suggesting that this approach was well-founded. This plot represents the power curves from this analysis alongside the position of each experiment in the manuscript that used a similar round-based structure in this landscape. Please note that we aimed for Experiments 3A and 3B to have adequate power to detect an effect in each experimental condition, and so this plot displays power per experimental condition (12 decisions each) rather than for the full experiment, and Experiment 4 is separated into main rounds (“4”) and the final between-subjects round (“4f”).

